# Changes in aperiodic (1/*f* slope) activity during a picture-word interference task: Effects of congruency and sequence manipulations

**DOI:** 10.64898/2025.12.19.695375

**Authors:** Virginia Tronelli, Patrycja Kałamała, Gabriele Gratton, Monica Fabiani, Mate Gyurkovics, Kathy A. Low, Maurizio Codispoti, Andrea De Cesarei

## Abstract

Aperiodic neural activity (1/*f* EEG) has been proposed to reflect the balance between excitatory and inhibitory (E/I) inputs, with steeper spectral slopes reflecting increased inhibition and flatter slopes indicating excitation. This activity is also thought to reflect the temporal coordination of neural firing, offering insights into fundamental brain dynamics. Recent studies have shown that the 1/*f* slope is sensitive to stimulus onset, characterized by initial inhibitory shifts followed by excitatory rebounds, which may reflect cognitive control mechanisms involved in suppressing distractions and preparing goal-directed responses. However, previous research has relied on fixed temporal windows and insufficient control of ERP contamination, limiting our understanding of rapid control dynamics. Here we used newly developed time-resolved analyses to study 1/*f* spectral slope modulation during a Picture-Word Interference task, focusing on two canonical cognitive control markers: the Congruency Effect (CE) and Congruency Sequence Effect (CSE). Forty-nine participants categorized pictures while ignoring congruent or incongruent words. Behaviorally, we replicated robust CE and CSE patterns. Spectral slope analyses showed that incongruent trials elicited steeper slopes — consistent with increased inhibition — particularly in frontal and central regions, reflecting conflict-related control engagement. Moreover, CSE analyses revealed dynamic slope modulations across frontal, central, and occipital components over time, suggesting control adjustments influenced by previous trial congruency. These results provide the first within-trial time-resolved evidence that aperiodic 1/*f* EEG activity can track both immediate conflict resolution and cognitive adjustments, offering a temporally sensitive neural marker of cognitive control, albeit with effects that are small in magnitude on average.

## 1. Introduction

Aperiodic neural activity (also called 1/*f* component due to its shape in the power spectrum) has gained increasing recognition as a potential marker of cortical Excitation-Inhibition (E/I) balance in electroencephalographic (EEG) research (Ahmad et al., 2022; Gao et al., 2017). Recently, cognitive neuroscientists have begun to link aperiodic activity and E/I balance with cognitive control mechanisms (Donoghue et al., 2020; Gyurkovics et al., 2022; Voytek et al., 2015; Waschke et al., 2021; see also Gratton et al., 2018). However, how aperiodic parameters evolve over time to support information processing remains unclear. In this study, we use time-resolved analyses to investigate the temporal evolution of aperiodic EEG during the congruency effect (CEs) and the congruency sequence effect (CSE) — two canonical markers of cognitive control (Gratton et al., 2018; von Bastian, 2020) — offering a novel perspective on the neurophysiological mechanisms underlying these phenomena.

### 1.1 Cognitive Control

Cognitive control enables the adaptive regulation of behavior in response to changing environmental demands (Gratton et al., 2018; von Bastian, 2020). A key behavioral marker of cognitive control is the CE—a robust phenomenon in conflict tasks whereby performance systematically depends on the congruency between task-relevant and task-irrelevant information (Eriksen & Eriksen, 1974; Gratton et al., 1992). Incongruent trials, which require resolving competing stimulus–response associations, typically elicit slower responses and more errors than congruent trials. The CE emerges from the interplay of distinct processes engaged during task performance, including perceptual, attentional, and executive mechanisms. Nevertheless, its most prominent interpretation is as an index of cognitive control: the interference captured by the CE represents the very conditions in which cognitive control must be engaged, i.e., a situation where task-irrelevant information impedes goal-directedness. Consequently, greater interference necessitates stronger top-down regulation, which is why the CE is widely regarded as a central marker of cognitive control (Eriksen & Eriksen, 1974; Gratton et al., 1992; von Bastian, 2020).

Beyond this immediate performance cost, cognitive control also shows adjustments based on trial history. This is reflected in the CSE (Gratton et al., 1992; Egner, 2007)—a reduction of the CE following incongruent relative to congruent trials, typically interpreted as a transient upregulation of cognitive control (Botvinick et al., 2001). The CSE has been explained by several theoretical accounts, with some proposing the engagement of cognitive control and others suggesting it operates without direct control involvement. The most prominent cognitive control account, the conflict-monitoring theory (Botvinick et al., 2001), posits a dedicated conflict-monitoring system that detects the simultaneous activation of competing response tendencies, as occurs in incongruent trials. The resulting conflict signal triggers an upregulation of cognitive control, which enhances performance on subsequent incongruent trials, reducing the CE. At the neural level, activity in the conflict-monitoring unit has been linked to the anterior cingulate cortex (ACC), while the upregulation of control is thought to rely on the dorsolateral prefrontal cortex (DLPFC; Botvinick et al., 2001). Alternative non-control related accounts emphasize the involvement of learning and memory processes rather than cognitive control. For instance, *episodic retrieval*, *feature integration*, and *contingency learning* accounts propose that trial-to-trial changes may be driven by automatic processes, such as retrieval of stimulus-response bindings, stimulus repetitions, or learned associations between stimuli and responses (for a review, see Braem et al., 2019). Importantly, research shows that the CSE can still occur in the absence of these factors (Jiménez & Méndez, 2014; Kim & Cho, 2014; Schmidt & Weissman, 2014; Weissman et al., 2014; Gyurkovics, Stafford, & Levita, 2020; Gyurkovics, Kovacs, et al., 2020; Gyurkovics & Levita, 2021). Overall, evidence indicates that the CSE likely reflects a combination of cognitive control and non-control mechanisms, making it a well-established albeit impure indicator of control adjustments (Abrahamse et al., 2016; Egner, 2023).

Both the CE and CSE have been extensively investigated as markers of cognitive control in behavioral and neurophysiological research (e.g., Botvinick et al., 2001; Cavanagh & Frank, 2014; Cohen & Donner, 2013; De Cesarei et al., 2023; Gratton et al., 1992; Gyurkovics & Levita, 2021; Schiltenwolf et al., 2024; Tronelli et al., 2025; van Maanen et al., 2009). To date, most of the EEG evidence about the neural underpinnings of CE and CSE has come from event-related potential (ERP) and time-frequency research, which primarily emphasize phase-locked or rhythmic brain activity associated with conflict resolution and cognitive adjustments, respectively. More recently, growing interest has emerged in the non-rhythmic, non-phase-locked aperiodic component of the EEG signal (1/*f*) as a complementary source of information about cognitive function (Gao et al., 2017).

### 1.2 1/*f* EEG activity

This aperiodic neural activity, also referred to as 1/*f* noise, is characterized in the frequency domain by a gradual decline in power with increasing frequency. This component is best quantified in log-log space, where its two key parameters—exponent and offset—can be extracted from the slope and intercept of the power spectrum after separating out periodic (oscillatory) activity. Specifically, the relationship can be described by the equation below:

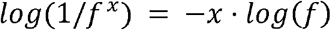

such that a steeper (more negative) slope, corresponding to a higher (more positive) exponent (*x* in the formula), reflects a stronger decay of power with increasing frequency (*f* in the formula). Conversely, a flatter (less negative) slope, corresponding to a lower (less positive) exponent, indicates a weaker decay.

At the level of neural mechanisms, aperiodic activity has been linked to the balance between excitatory and inhibitory inputs in neural circuits (called E/I balance, Gao et al., 2017). Within this framework, steeper spectral slopes are thought to reflect increased inhibitory activity and flatter spectral slopes indicate a relative increase in excitatory activity, suggesting that properties of aperiodic activity may provide insight into fundamental aspects of brain functioning (Ahmad et al., 2022; Donoghue et al., 2020; Gao et al., 2017; Gyurkovics et al., 2022; Voytek et al., 2015; Waschke et al., 2021). Relatedly, the spectral slope is also theorized to serve as an index of the temporal coordination of neural firing: steeper slopes suggest more coordinated, less noisy signaling, while flatter slopes are associated with noisier neural activity (Chini et al., 2022; He, 2014; He et al., 2019; Voytek & Knight, 2015). These interpretations align with neurophysiological models of cognitive control, which propose that low-frequency power reflects feedback loops involving inhibitory connections, while high-frequency power reflects feedforward, excitatory activity, supporting both auxiliary and task-relevant processes, not necessarily limited to cognitive control (Gratton, 2018). Within this framework, stimuli, task demands, or contextual variability can transiently recruit inhibitory mechanisms that support cognitive performance, producing steeper spectral slopes.

Gyurkovics et al. (2022) were among the first to demonstrate systematic stimulus-induced changes in the aperiodic EEG. They reported more negative post-stimulus spectral slopes (compared to the pre-stimulus interval) that were independent of concurrent ERPs and scaled with the cognitive demands of an auditory task. This pattern has been interpreted as a stimulus-driven shift in E/I balance toward increased inhibition, potentially reflecting a transient suppression of ongoing excitatory activity. Such a mechanism may facilitate the reallocation of neural resources and the flexible updating of mental representations in response to task-relevant stimuli, which is a primary function of cognitive control.

Subsequent studies have provided further evidence that aperiodic EEG can be modulated by experimental manipulations (e.g., Akbarian et al., 2024; Frelih et al., 2024; Jia et al., 2024; Kałamała et al., 2024; Lu et al., 2024; Manyukhina et al., 2024; Yan et al., 2024; Zhang et al., 2023). For example, Kałamała et al. (2024) provided the first evidence of *within-trial* modulations in aperiodic activity. By segmenting each trial into 500-ms intervals, the authors showed that the presentation of a stimulus (a cue in a cued flanker task) initially produced a more negative spectral slope compared to the pre-cue period (as found in Gyurkovics et al., 2022), which progressively transitioned into a more positive slope as time passed. They interpreted this pattern as reflecting an initial increase in inhibition, likely serving to suppress ongoing neural activity and facilitate the processing of task-relevant information, followed by a rise in excitation in support of motor response preparation for the upcoming imperative stimulus. These mechanisms are all associated with cognitive control, which fundamentally aims to ensure goal-directedness in the face of distractions (Gratton et al., 2018; von Bastian et al., 2020). Relatedly, recent studies have demonstrated *across-trial* variability in the aperiodic component (Boudewyn et al., 2025; Jia et al., 2024). For instance, Jia et al. (2024) showed that the aperiodic EEG component tracks changes in the CE across adjacent trials in a flanker task. Specifically, spectral slopes were more negative in the incongruent condition compared to the congruent condition during the current trial, but this pattern reversed in the subsequent trial, with more negative slopes observed in the congruent condition than in the incongruent condition. This reversal suggests that the post-stimulus spectral slope is not simply a continuation of pre-stimulus activity but instead indicates a shift in neural dynamics across trials.

While existing studies offer promising evidence that stimulus-induced changes in the aperiodic component of the EEG may reflect cognitive control mechanisms, their conclusions are limited by several methodological shortcomings. Some of them rely on simple paradigms with limited performance measures (e.g., Gyurkovics et al., 2022), making it difficult to assess the behavioral relevance of the observed spectral slope changes. Moreover, most studies do not account for ERPs (e.g., Jia et al., 2024; Zhang et al., 2023), which, as transient, non-oscillatory patterns of activity, can confound the estimation of aperiodic parameters (for a detailed argument, see Gyurkovics et al., 2022). Critically, these analyses often rely on averaging neural responses over broad, fixed time windows (typically longer than 500 ms), which may obscure the more rapid temporal dynamics of aperiodic activity. This last limitation is particularly important because cognitive control is a transient, adaptive process that unfolds rapidly following conflict-related information (Gratton et al., 1992, 2018; Botvinick et al., 2001). Capturing moment-to-moment fluctuations in aperiodic activity is therefore essential for understanding how cognitive control is dynamically deployed and adjusted over time. While Kałamała et al. (2024) challenged the notion that the aperiodic signal remains stable throughout a task trial, no prior study has tracked time-resolved changes in aperiodic activity in response to canonical manipulations of cognitive control, such as the CE and CSE (but see Frelih et al., 2024, for a recent time-resolved analysis in an n-back task).

### 1.3 The current study

In this study, we investigated the aperiodic characteristics of post-stimulus EEG related to the CE and CSE using a within-trial, time-resolved approach, which allows us to track dynamic changes unfolding within a single trial, in contrast to trial-by-trial analyses that capture variability across trials. To achieve this, we analyzed scalp EEG from young adults performing a Picture-Word Interference (PWI) task (Rosinski, 1977; Tronelli et al., 2025), in which participants are asked to categorize pictures as either animals or vehicles while ignoring a superimposed word. The word can either be congruent (e.g., a picture of an animal with the word "animal") or incongruent (e.g., a picture of an animal with the word "vehicle"), manipulating conflict level and cognitive control engagement.

To capture the temporal dynamics of aperiodic activity, stimulus-locked epochs underwent time-frequency decomposition using wavelet convolution, and the spectral slopes were estimated from the resulting power spectral densities time point by time point using a censored regression-based approach (Kałamała et al., 2025). Single-cycle wavelets were selected to prioritize temporal over frequency resolution and thereby increase sensitivity to brief, transient changes. The resulting estimates are time-resolved rather than truly instantaneous: power at each nominal time point is integrated over the finite temporal support of the corresponding wavelet, which is longer at lower than at higher frequencies (for details, see *Methods*). Consequently, each slope combines power estimates with frequency-dependent temporal support and is not equivalent to a conventional aperiodic slope calculated within a single fixed window applied uniformly across frequencies. This approach was used specifically to examine rapid within-trial changes that would be attenuated or temporally blurred by the relatively long fixed windows required to estimate power at the lowest frequency included in the analysis.

We expected that both the CE and CSE would be reflected in changes in aperiodic EEG activity. Building on theories that attribute CE and CSE to the engagement of cognitive control (Botvinick et al., 2001; Gratton et al., 1992), as well as on research linking steeper slopes to higher cognitive demands (Gyurkovics et al., 2022; Kałamała et al., 2024; Lu et al., 2024), potentially through increased cortical inhibition, we hypothesized that incongruent trials would be associated with more negative spectral slopes after stimulus presentation compared to congruent trials. Furthermore, we expected that the preceding trial would modulate the E/I ratio, as captured by the spectral steepness of the aperiodic activity in the current trial. Specifically, we predicted that the difference in spectral slope steepness between current incongruent and congruent trials would be greater following congruent trials than following incongruent trials, mirroring the behavioral CSE pattern. At the statistical level, we used Principal Component Analysis (PCA) to reduce spatial variability (spatial PCA) and to infer the effective temporal window for the analyses (temporal PCA). Because this study introduces a novel approach to aperiodic slope estimation—based on single-cycle wavelet convolution—we report two complementary statistical analyses: (i) an exploratory repeated-measures ANOVA, conducted on spatial PCA components within time windows derived from the temporal PCA, without correction for multiple comparisons; and (ii) cluster-based permutation analyses^1^ applied to the same spatial PCA components, controlling for multiple comparisons at the family-wise error rate (FWER) level. While the first analysis provides exploratory leads for future work, the second allows us to assess which effects are robust.

By tracking the moment-to-moment evolution of the spectral slope, we aimed to reveal how E/I balance shifts over time, translating into conflict resolution and cognitive control adjustments. This approach offers a novel perspective on the neural mechanisms underlying CE and CSE, potentially uncovering transient shifts in cortical E/I balance that contribute to flexible cognitive control.

## 2. Method

### 2.1 Participants

The study was conducted at the University of Bologna, Italy. Fifty Italian-speaking participants with normal or corrected vision took part in the experiment. Data from 49 participants (20 females, mean age ± *SD* = 26.27 ± 3.38) were analyzed due to one participant failing to comply with task instructions. The experimental protocol adhered to the principles outlined in the Declaration of Helsinki and received approval from the Ethical Committee of the University of Bologna. Written informed consent was obtained from all participants.

### 2.2 Stimuli

The Picture Word Interference (PWI) task involved a set of 488 different pictures sourced from the internet, which were employed in previous studies investigating visual categorization and cognitive control (De Cesarei et al., 2019, 2021, 2023; Tronelli et al., 2025). Each picture depicted either one or two animals or vehicles in an indoor or outdoor setting, resulting in eight equally probable combinations across three orthogonal dimensions: content (animal vs. vehicle), number of foreground elements (one vs. two), and scenario (indoors vs. outdoors). To ensure consistency, all pictures met the following criteria: only one or two clearly visible animals or vehicles in the foreground, no additional animals or vehicles in the background, and a clearly distinguishable setting.

The pictures were presented in two blocks: one block in which pictures did not repeat across trials (i.e., Novel Picture Block) and the other block in which the same pictures were repeatedly shown across trials (i.e., Frequent Picture Block). In the Novel Picture Block, 480 pictures out of the total 488 were used. In the Frequent Picture Block, the remaining eight pictures were used to create four pairs of pictures, each consisting of an animal and a vehicle, and during the presentation of Frequent Picture Block participants saw one of the four repeated picture pairs. Hence, participants viewed 480 pictures, all different in the Novel Picture Block, and a pair of pictures out of the four repeated 240 times in the Frequent Picture Block. In the Frequent Picture Block, the pictures were repeated across trials so that response repetition to the picture category (i.e., animal or vehicle) from one trial to another also implied the repetition of the same picture. In the Novel Picture Block, all the pictures were different, so the repetition of the response to the picture category only implied the repetition of the same response category (animal or vehicle), as shown in **Figure 1**.

**Figure 1.**
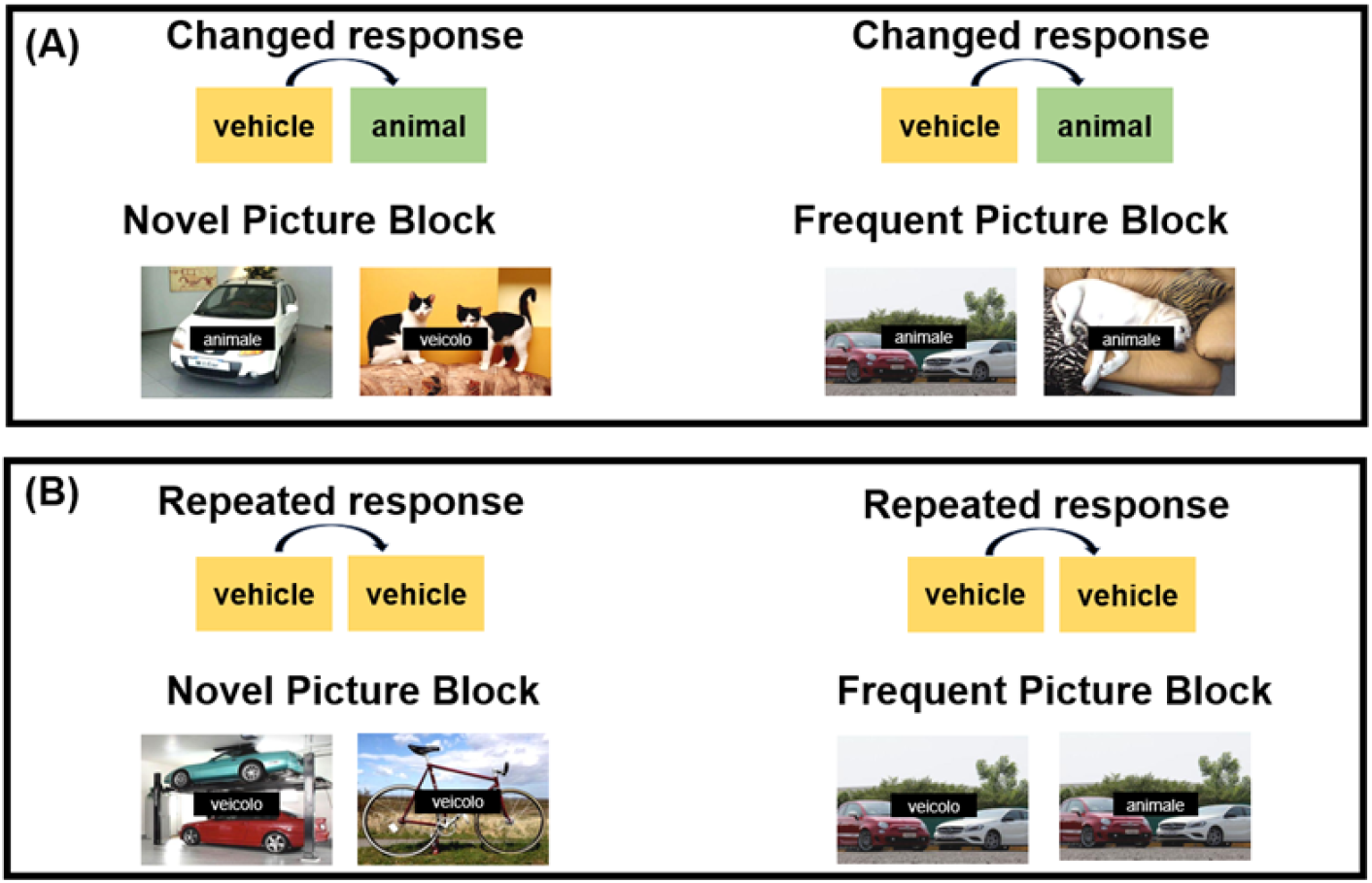
Overview of experimental design. Examples of two consecutive trials in the changed response category condition (Panel A) and in the repeated response category condition (Panel B), shown separately for the Novel Picture Condition (left panels) and the Frequent Picture Condition (right panels). The distractor word could be congruent with the picture (e.g., the word animale — animal in English — on an image of an animal) or incongruent (e.g., the word veicolo — vehicle in English — on an image of an animal).

The pictures were displayed in color, with brightness and contrast adjusted to ensure consistency. The average pixel intensity was 153 (*SD* = 5.05) on a scale from 0 to 255. Each full-screen picture was resized to a 1280 × 1024 pixel monitor, subtending 20°30’ horizontal × 16°20’ vertical degrees of visual angle. Each picture was shown together with a word that was congruent or incongruent with the content of the picture. The words were in Italian: "veicolo" ("vehicle") and "animale" ("animal") (e.g., a picture that represents a cat with the word "animale" - i.e., "animal" - is a congruent trial, while the same picture with the word "veicolo" – i.e., "vehicle" - is an incongruent trial). The white words in Courier New font (size 70) were centered on the screen within a black rectangle (15°8’ horizontally × 2°33’ vertically) superimposed on the picture.

### 2.3 Procedure

During the PWI task, participants were seated in the experimental room, where the illumination was set to 3 lux, as measured with a diode-type digital luxmeter. The experiment comprised two blocks of 480 trials: the Novel Picture Block (480 trials, each featuring a unique picture) and the Frequent Picture Block (480 trials, in which two pictures were shown 240 times each), totalling 960 trials. Participants were instructed to respond to the picture while ignoring the word, giving equal importance to response speed and accuracy. They pressed one of two keys (J or N) on the computer keyboard using two fingers of their dominant hand. The order of task blocks and response keys were counterbalanced across participants. The experiment lasted approximately 60 min, with eight practice trials preceding each experimental block to familiarize participants with the task. The distance between the monitor and the participant was 94 cm.

Each trial began with a 500-ms picture presentation, followed by a response-stimulus interval (RSI) of either 1000, 2000, 3000, or 5000 ms. Out of the 960 trials, half were congruent and half incongruent. The sequence of trials was designed such that each of the four possible sequences (e.g., previous trial congruent or incongruent, followed by a congruent or incongruent trial) occurred with equal probability (120 trials per sequence per block). Additionally, each sequence was equally paired with each RSI condition, resulting in 30 trials per condition in each block. In summary, the PWI task comprised five experimental manipulations: Stimulus Repetition (Novel Picture Block vs. Frequent Picture Block), Category-Response Repetition (Same Category vs. Changed Category), RSI (1000, 2000, 3000, or 5000 ms), Previous Congruency (Previous-Congruent vs. Previous-Incongruent), and Current Congruency (Congruent vs. Incongruent). Since the present study focuses specifically on the well-established CE and CSE, we limited our analysis to these two manipulations. All other conditions were collapsed, as they fall outside the scope of the current research. The experiment was run using E-Prime 2.0 Professional.

### 2.4 Behavioral Analysis

The dependent variables were mean accuracy and mean reaction time (RT). Practice trials and the first trial of each block were excluded. For the RT analysis, we additionally excluded trials with incorrect responses, trials following an error, and trials with RTs exceeding ±2.5 standard deviations (*SD*s) from the participant’s mean. Both accuracy and RT data were analyzed with a repeated-measures ANOVA, with the following within-subject factors: Previous Congruency (two levels: Previous-Congruent, Previous-Incongruent) and Current Congruency (two levels: Congruent, Incongruent). Huynh–Feldt correction for lack of sphericity was used when appropriate. Follow-up paired *t*-tests were performed in the case of significant interactions. A *p*-value < .05 was considered the threshold for statistical significance. For all tests, partial eta squared (η*_p_²)* and Cohen’s d were calculated and reported as measures of effect size for ANOVAs and t-tests, respectively.

### 2.5 EEG Data Acquisition and Preprocessing

Scalp EEG was recorded from 64 Ag/AgCl active electrodes using a BioSemi ActiveTwo^®^ system (Amsterdam, The Netherlands). The electrodes were secured by an elastic cap according to the extended 10-20 international electrode placement system (Jasper, 1958). Horizontal and vertical electrooculograms (EOGs) were also recorded to monitor ocular artifacts. The sampling rate was 512 Hz. During recording, data were referenced to the common mode sense (CMS) electrode and were filtered online with a low pass filter equal to 1/5 of the sampling rate (i.e., 102.4 Hz).

The data were pre-processed using custom MATLAB 2022b codes (The MathWorks) incorporating EEGLAB 13.6.5 (Delorme & Makeig, 2004) and ERPlab 6.1.3 (Lopez-Calderon & Luck, 2014). The EEG was first re-referenced to the average mastoids and bandpass filtered with 0.1 and 40 Hz cut-off frequencies. The data were then segmented into 2000-ms epochs, spanning from 500 ms before to 1500 ms after stimulus onset. After excluding epochs with amplifier saturation and performing ocular correction (Gratton et al., 1983), epochs with peak-to-peak voltage fluctuations at any EEG electrode exceeding 150 μV (600-ms window width, 100-ms window step) were discarded. Additional epochs were excluded if they contained an incorrect response, were followed by an error, were the first trial of a block, or had a RT exceeding ±2.5 *SD*s from the participant’s mean to match the RT analysis criteria. Data from electrodes Fp1, Fp2, AF3, AF4, AF7, AF8, AFz, and FPz were excluded as they often contain minor residual ocular artifacts even after ocular correction. This left 56 electrodes for further analysis. The average number of artifact-free epochs per participant was 694 (*SD* = 153, *min* = 366, *max* = 893).

### 2.6 Spectral Analysis

To examine the temporal variation of the aperiodic EEG component, stimulus-locked epochs underwent wavelet decomposition, and the spectral slopes were estimated from the resulting power spectral densities at each time point using a censored regression-based approach, as described below (see also Kałamała et al., 2025). For a description of the spectral analysis workflow see **Figure 2**. Data and code necessary to reproduce the statistical analyses (section 3.2.3) will be available at https://osf.io/z3qsg (view only link for reviewers: https://osf.io/z3qsg/overview?view_only=fc282671ba704386bf47f7776d4e4509).

**Figure 2.**
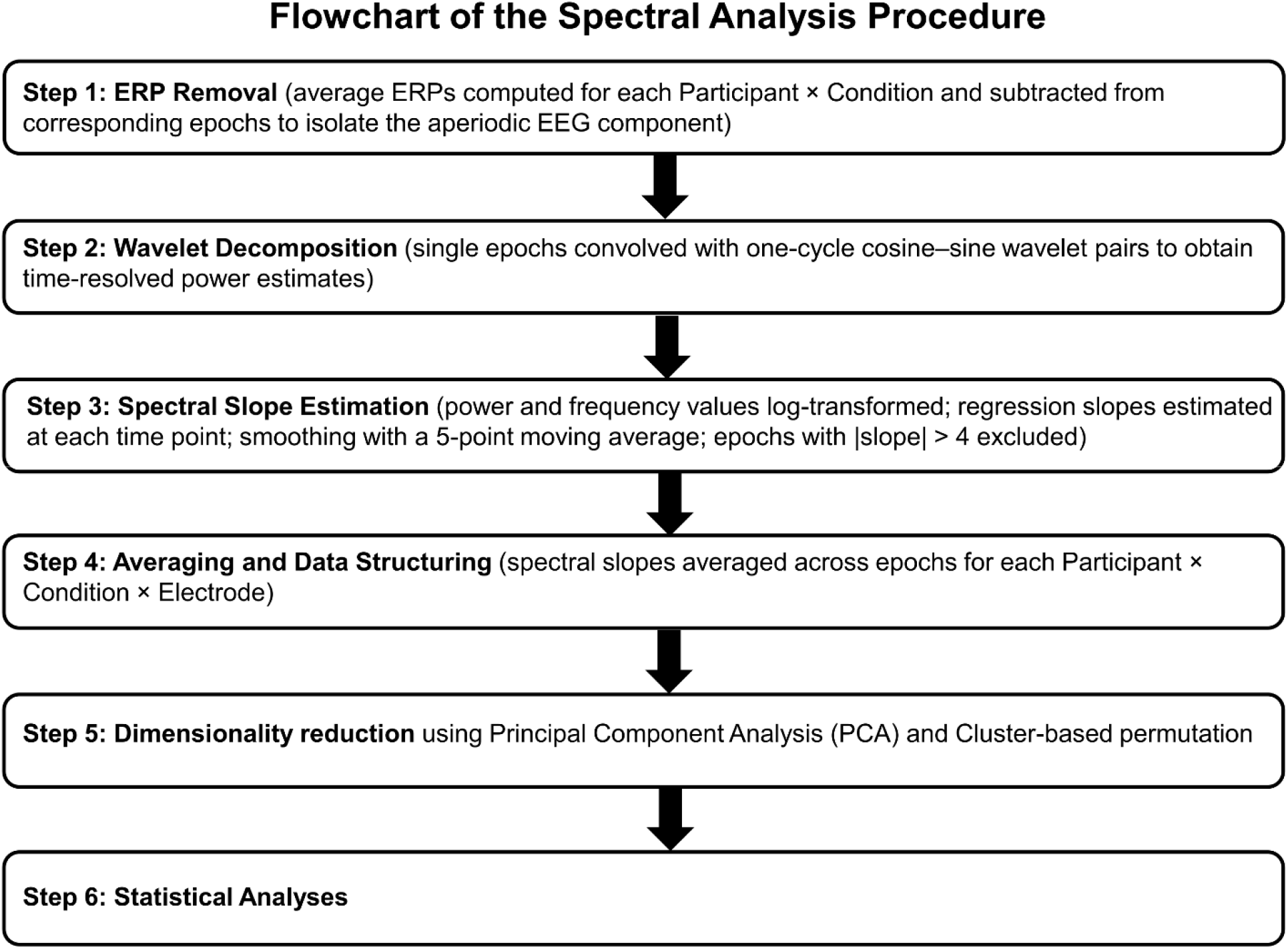
Flowchart of the Spectral Analysis Procedure. The diagram shows the main processing steps.

First, we removed the average event-related potentials (ERPs) from the EEG data to avoid distortions in aperiodic parameter estimation. Since ERPs contribute to the overall EEG spectrum in a broadband fashion, removing them allows for a more accurate isolation of the aperiodic signal (for further discussion, see Gyurkovics et al., 2022). This was done by computing the average ERP for each Participant × Condition combination and subtracting it from the corresponding epochs, following established procedures (Gyurkovics et al., 2022; Kałamała et al., 2024).

Next, the single epochs were convolved with sliding cosinusoidal and sinusoidal wavelet pairs, which were combined to provide time-resolved power estimates for each frequency. These estimates are not truly instantaneous: each value summarizes signal information over the finite temporal support of the corresponding wavelet, which spans one cycle and therefore differs across frequencies. The temporal support ranged from approximately 400 ms at 2.5 Hz to 40 ms at 25 Hz. Consequently, the slope estimated at a given nominal time point combines power estimates reflecting different temporal integration intervals. One-cycle wavelets were used to prioritize temporal over frequency resolution and thereby increase sensitivity to brief, rapidly evolving changes in aperiodic activity. Wavelets were not weighted with Gaussian tapers and simply constituted a single cycle of a sine or a cosine wave at a given frequency. The analysis focused on a limited set of frequencies to optimize computational time and minimize redundancy due to overlap between nearby frequencies. Because the emphasis here was on *temporal resolution*, this necessarily limited the frequency resolution of the analyses. This was considered acceptable since the focus is on broadband rather than narrowband activity. Specifically, we selected values corresponding to successive powers of 2 within the available frequency range—i.e., the 2nd, 4th, 8th, 16th, and 32nd frequencies from the set—resulting in the following values: 2.5, 5, 7.5, 15, and 25 Hz. Restricting the frequency range facilitates more accurate estimation of the spectral slope by excluding frequencies most likely to reflect periodic activity. Prior research has shown that most scalp-recorded oscillatory activity during wakefulness falls within the theta-alpha-beta range (e.g., Myrov et al., 2024). Accordingly, power values within the 7.5–15 Hz range were excluded from slope and intercept estimation. This procedure yields estimates that more accurately capture the aperiodic component.

The resulting power estimates and corresponding frequencies were log-transformed, and regression slopes were calculated for each time point using a least-squares approach to provide time-resolved estimates of the aperiodic component. These estimates should not be considered equivalent to slopes calculated from a conventional fixed temporal window applied uniformly across frequencies. Rather, they were designed to prioritize sensitivity to rapid changes in the broadband spectral profile. To temporally smooth the data, slope estimates were filtered using a 5-point moving average within each epoch. Epochs with slope values exceeding ±4 were considered outliers and excluded. On average, 659 epochs per participant were retained (*SD* = 168, *min* = 274, *max* = 891). The remaining slopes were then averaged across epochs for each EEG electrode (56 electrodes in total) and experimental condition (Stimulus Repetition × Category Repetition × RSI × Previous Congruency × Current Congruency) within each participant.

#### Parametrization and data reduction

To reduce data dimensionality, improve the signal-to-noise ratio, and increase statistical power, the Participant × Electrode × Condition × Time data were subjected to Principal Component Analysis (PCA). Because the wavelet decomposition introduced edge artifacts at the beginning and end of each epoch, the first and last 200 ms were excluded prior to PCA. Thus, all PCA analyses were conducted on the interval from −300 to 1300 ms relative to stimulus onset.

The PCA procedure involved two stages: a spatial PCA followed by a temporal PCA. In the first stage, PCA was applied to the electrode dimension (spatial PCA). For this analysis, the data matrix was arranged such that observations corresponded to Participant × Condition × Time, and variables corresponded to EEG electrodes. This decomposition identified spatial components reflecting patterns of covariance across electrodes. Once the spatial components were extracted, the EEG data were projected onto the spatial loadings using the regression method (i.e., EEG data × unstandardized factor loadings × inverse covariance matrix; Thomson, 1938; Thurstone, 1935), yielding spatial factor scores organized as Participant × Spatial Component × Condition × Time.

In the second stage, PCA was applied to the time dimension (temporal PCA). For this analysis, the data matrix was reorganized so that observations corresponded to Participant × Condition × Spatial Factor Scores, and variables corresponded to time points within the epoch. In other words, the temporal PCA was conducted on the spatial factor scores derived from the first-stage spatial PCA. This decomposition identified temporal components reflecting patterns of covariance across time and was used to characterize the dominant temporal structure of the signal.

In both the spatial and temporal PCA, the number of components to retain was first determined using the Empirical Kaiser Criterion (EKC; Braeken & Van Assen, 2017; Steiner & Grieder, 2020). Specifically, a correlation matrix was computed for the variables entered into a given decomposition (electrodes for the spatial PCA; time points for the temporal PCA), and the EKC was used to identify components explaining more variance than expected from random noise. Once the number of retained components had been established, an initial unrotated PCA solution was estimated from the covariance matrix using a factor-analytic routine, yielding unstandardized factor loadings. These loadings describe the structure of each component (spatial or temporal), whereas factor scores are a weighted combination of observations that quantify the expression of each component across said observations. To improve interpretability, the retained components were then subjected to varimax rotation (Kaiser, 1958). This orthogonal rotation redistributes variance across components to achieve a simpler and more interpretable loading structure by emphasizing high-contrast loadings. After rotation, any component explaining less than 1/x of the total variance (where x is the total number of retained components) was excluded from further analysis.

The temporal PCA was used to establish effective temporal resolution for subsequent statistical analyses, i.e., to define the optimal time window within which effects can be reliably detected. To summarize the typical temporal extent of these components, we calculated the Full Width at Half Maximum (FWHM) for each temporal component, given their roughly Gaussian shape (see Section 3.2.2 for further details). This estimate was then used to divide the −300 to 1300 ms interval into consecutive time windows anchored to stimulus onset (0 ms). Importantly, these windows were not intended to correspond exactly to individual temporal components, but rather to approximate the characteristic temporal scale of the structured variance identified by the temporal PCA.

In summary, the spatial PCA reduced the dimensionality of the electrode space and yielded spatial components for further analysis, whereas the temporal PCA, applied to the spatial component scores, was used to estimate the characteristic temporal scale of the data and thereby define the time windows used in the statistical analyses.

#### Statistical testing

We performed two complementary statistical analyses: (i) an exploratory repeated-measures ANOVA and paired *t*-tests conducted on the spatial PCA components within time windows defined by the temporal PCA, without correction for multiple comparisons; and (ii) time-resolved cluster-based permutation analyses applied to the same spatial PCA components, controlling for multiple comparisons at the family-wise error rate (FWER) level. Analyses were conducted separately for each spatial component using the corresponding spatial scores (derived from the spatial PCA). For the exploratory analyses, scores were averaged within each FWHM-defined time window (as determined by temporal PCA), whereas the permutation analyses were performed across time points.

#### Exploratory analyses (no correction for multiple comparisons)

Given that examining the effects of experimental manipulation on aperiodic activity in a time-resolved manner is a novel approach in EEG research, we first assessed the global stimulus-induced changes, independent of experimental condition. This was done by comparing condition-averaged values in each post-stimulus PCA-based time window to the pre-stimulus window using paired *t*-tests. The PCA, as well as the spectral decomposition, were performed across all available conditions and then averaged, so that only Previous and Current Congruency were included in the analyses. Subsequently, we conducted a repeated-measures ANOVA with Previous Congruency and Current Congruency as within-subject factors to evaluate the expected experimental effects, specifically the congruency and congruency sequence effects. Follow-up paired *t*-tests were performed in the case of significant interactions. Baseline correction was not applied, as no significant effects were observed in the pre-stimulus time window. A *p*-value < .05 was considered the threshold for statistical significance, without correction for multiple comparisons. For all tests, partial eta squared (η*_p_²*) and Cohen’s *d* were calculated and reported as measures of effect size for ANOVAs and *t*-tests, respectively.

### Cluster-based permutation analyses (FWER-corrected)

To complement the window-based exploratory analyses, we performed time-resolved cluster-based permutation tests on the same spatial PCA components. Analyses were conducted separately for each spatial component using the corresponding spatial scores and evaluated effects continuously across time, without predefined windows. Global post-stimulus changes were assessed relative to a pre-stimulus baseline, defined as the mean slope for *t* < 0 ms. Experimental effects were examined using within-subject contrasts corresponding to the 2 × 2 design: Congruency (Congruent vs. Incongruent), Previous Congruency (Previous-Congruent vs. Previous-Incongruent), and their interaction, defined as the difference in the Congruency effect as a function of Previous Congruency.

Statistical inference was based on a nonparametric sign-flip procedure (1,000 permutations), in which the sign of the data of a random half of participants was inverted on each permutation to generate a null distribution at each time point. Observed values were standardized relative to this null distribution to obtain *Z*-statistics (*Z*(*t*)). Clusters were defined as contiguous time points for which *Z*≥1.6449 or *Z*≤−1.6449, with positive and negative clusters identified separately. The cluster statistic was defined as the maximum absolute *Z*-value within each cluster. Statistical significance was determined by comparing each observed cluster statistic with the permutation-derived distribution of the maximum absolute cluster statistic across time and both directions, thereby controlling the two-sided FWER. Significant effects were defined as clusters with a permutation-based, two-sided FWER-corrected *p*<.05.

## 3 Results

### 3.1 Behavioral Data

#### 3.1.1 Accuracy

The main effect of Current Congruency was significant, *F*(1,48) = 30.24, *p* < .001, η*_p_ ^2^* = .39, with lower accuracy for incongruent (*M* = 96.74%, *SD* = 3.15) compared to the congruent trials (*M* = 97.53%, *SD* = 2.68), indicating the CE. The main effect of Previous Congruency was also significant, *F*(1,48) =6.80, *p* = .012, η*_p_ ^2^* = .12, with higher accuracy in trials following incongruent trials (*M* = 97.33%, *SD* = 2.73) compared to trials following congruent trials (*M* = 96.93%, *SD* = 3.14).

Finally, the two-way interaction between Current and Previous Congruency was significant, *F*(1, 48) = 12.02, *p* = .001, η*_p_²* = .20. There was no significant difference in accuracy between congruent trials preceded by congruent trials (*M* = 97.63%, *SD* = 2.64) and those preceded by incongruent trials (*M* = 97.43%, *SD* = 2.94), *t*(48) = 0.93, *p* = .357, *d* = .13. In contrast, the accuracy on incongruent trials was significantly higher when they were preceded by incongruent trials (*M* = 97.24%, *SD* = 2.67) compared to when they were preceded by congruent trials (*M* = 96.23%, *SD* = 3.79), *t*(48) = 4.12, *p* < .001, *d* = .59. When trials were preceded by a congruent trial, the accuracy was lower in incongruent trials compared to congruent trials, *t*(48) = −5.37*, p* < .001, *d* = -.77. There was no significant difference between incongruent trials and congruent trials when they were preceded by an incongruent trial, *t*(48) = -.98*, p* = .332, *d* = -.14 (indicating the CSE).

#### 3.1.2 Reaction Times

The main effect of Current Congruency was significant, *F*(1,48) = 16.68, *p* < .001, η*_p_ ^2^* = .26, with slower responses for incongruent (*M* = 656.39 ms, *SD* = 147.35) compared to the congruent trials (*M* = 641.84 ms, *SD* = 132.27), indicating the CE. The main effect of Previous Congruency was not significant, *F*(1,48) =.12, *p* = .732, η*_p_ ^2^* <.001.

The two-way interaction between Current and Previous Congruency was significant, *F*(1,48) = 18.33, *p* < .001, η*_p_ ^2^* = .28 (see **Figure 3**). Responses to incongruent trials were slower when they were preceded by a congruent compared with an incongruent trial, and responses to congruent trials were slower when they were preceded by an incongruent compared with a congruent trial, *t*(48) = 3.52, *p* =.001, *d* = .50 and *t*(48) = 2.87, *p* =.006, *d* = .41; respectively. There was no significant difference between incongruent and congruent trials when they were preceded by incongruent trials, *t*(48) = 1.44, *p* =.158, *d* = .21. Responses were slower during incongruent trials than during congruent trials when they were preceded by congruent trials, *t*(48) = 5.18, *p* <.001, *d* = .74 indicating the CSE.

**Figure 3.**
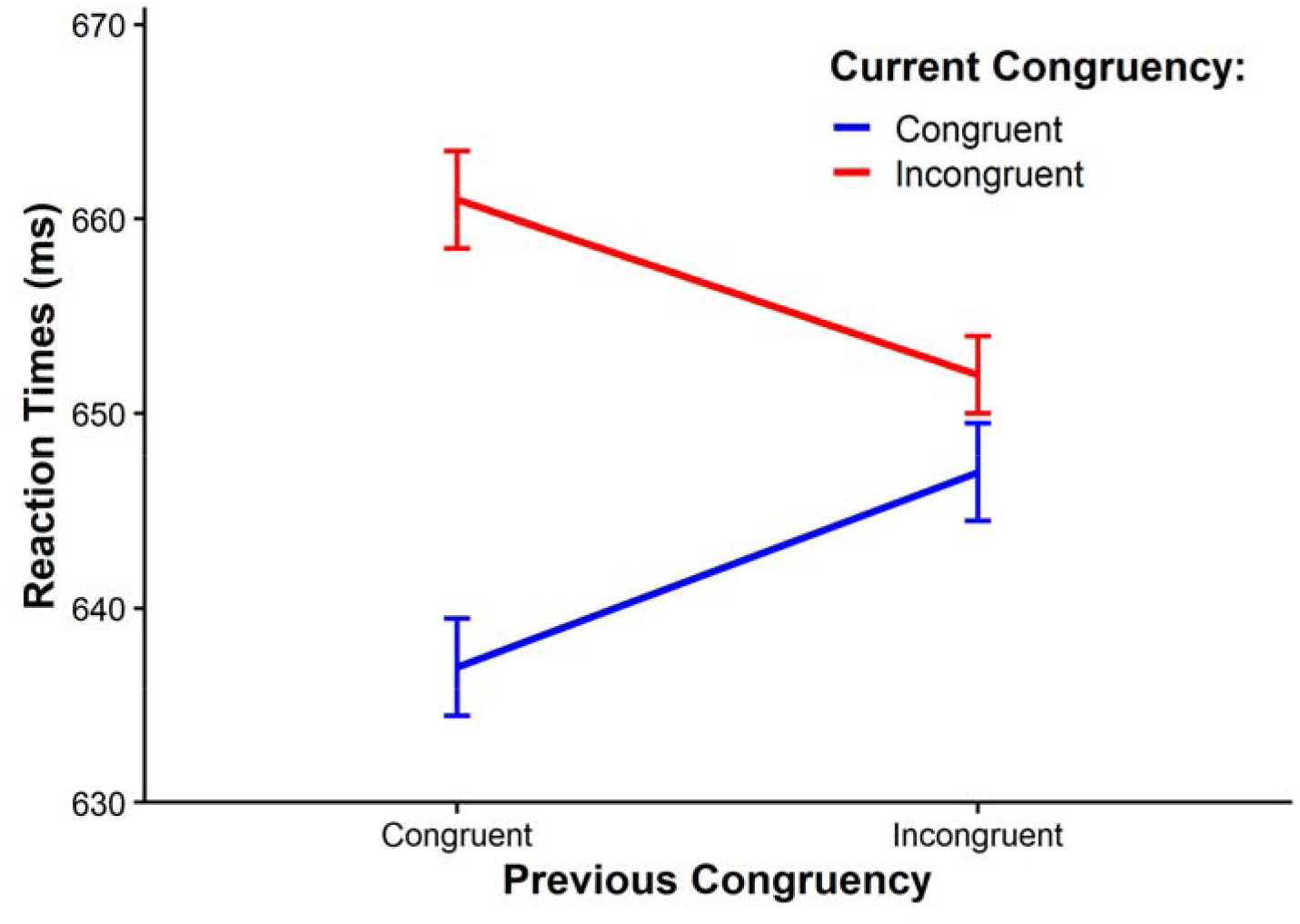
Current Congruency as a Function of Previous Congruency for Reaction Times. Mean reaction times for current congruency, broken down by previous congruency. Error bars represent ±1 within-subject standard errors of the mean (Cousineau, 2005).

### 3.2 EEG Data

#### 3.2.1 Spatial PCA

The Empirical Kaiser Criterion suggested a five-component solution. **Figure 4A** illustrates the distribution of loadings across electrodes, with each component showing a prominent peak in different scalp regions. Components were named after the region where they showed the maximum loading: occipital, frontal, central, left temporal, and right temporal. Temporally located components were excluded from further analysis as they accounted for less than one-fifth of the total variance (see **Figure 4B**). As a result, statistical analyses focused on the occipital, frontal, and central components.

**Figure 4.**
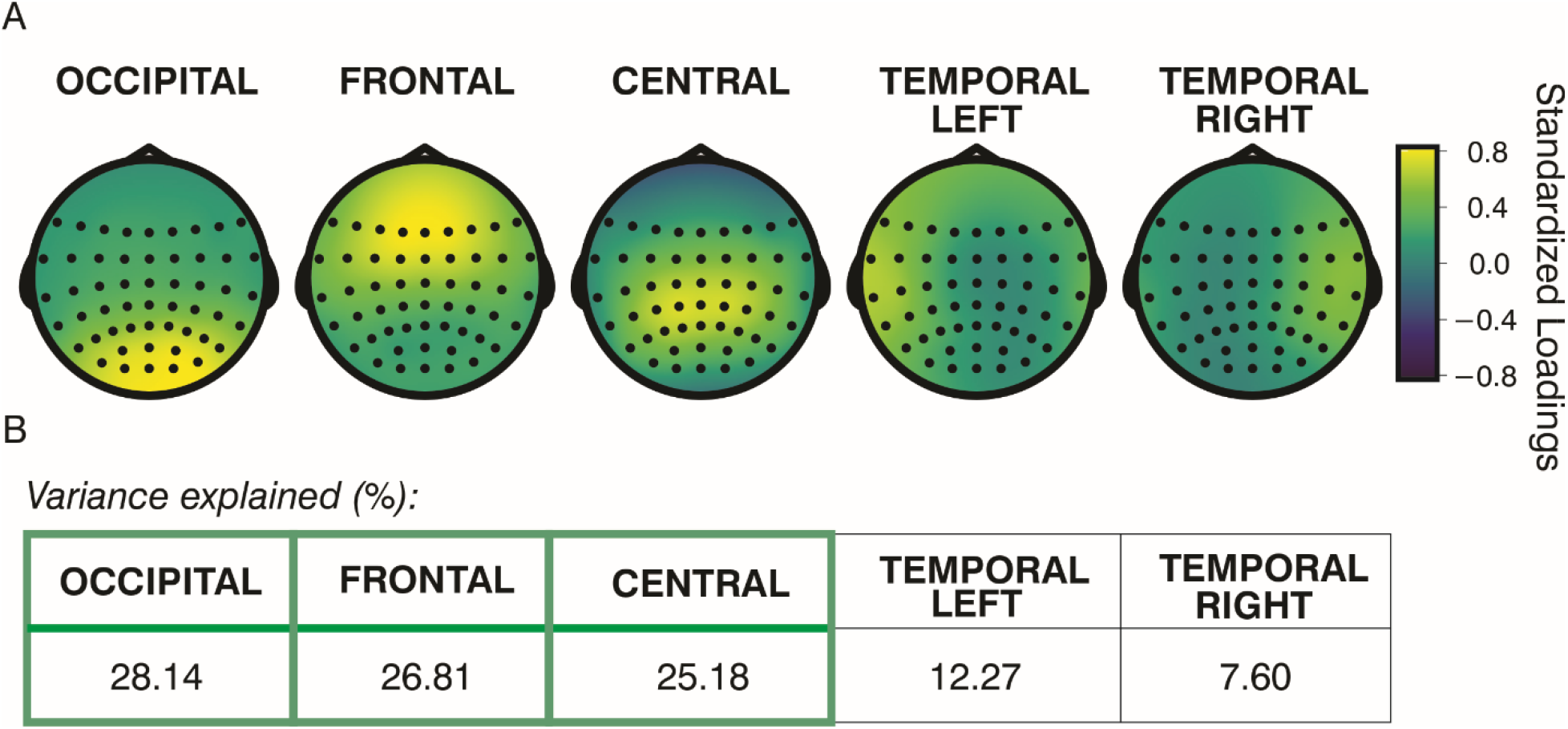
Results of the Spatial Principal Component Analysis. Distribution of electrode-wise standardized loadings after varimax rotation (Panel A), and percentage of variance explained by each corresponding component (Panel B). Components are ordered from highest to lowest explained variance. Those outlined in green in Panel B accounted for more than one-fifth of the variance and were retained for further analysis.

#### 3.2.2 Temporal PCA

The Empirical Kaiser Criterion indicated a 13-component solution. **Figure 5A** shows the distribution of loadings across time points, with each component showing a prominent peak in a different period. The resulting temporal loadings showed smooth, Gaussian-shaped profiles. This pattern is expected for the temporal PCA applied to EEG data, given the strong temporal autocorrelation of the signal and the variance-maximizing nature of the decomposition, which tends to produce smooth and temporally extended components. In addition, the varimax rotation may further accentuate the apparent temporal localization of components, yielding more peak-centered (Gaussian-like) profiles by concentrating variance within components and thereby enhancing interpretability. Five components were excluded as they accounted for less than 1/13 of the total variance (see **Figure 5B**). The mean FWHM across the remaining eight components was 162 ms (*Mdn* = 161 ms, *SD* = 25 ms). Based on this, a 160-ms period was selected as the effective time window for statistical analysis. Accordingly, the following time windows were defined: [-160,0], (0,160], (160,320], (320,480], (480,640], (640,800], (800,960], (960,1120], and (1120,1280], each containing 8 to 9 data points. Square brackets indicate the inclusion of the boundary value, while parentheses indicate exclusion. Time points at the edges of the epochs, where a full 160-ms time window could not be created, were excluded. For statistical testing, spectral slope values were averaged within each time window separately for each participant, condition, and spatial PCA component, yielding one value per window that was used as the dependent variable in the exploratory ANOVA and *t*-tests.

**Figure 5.**
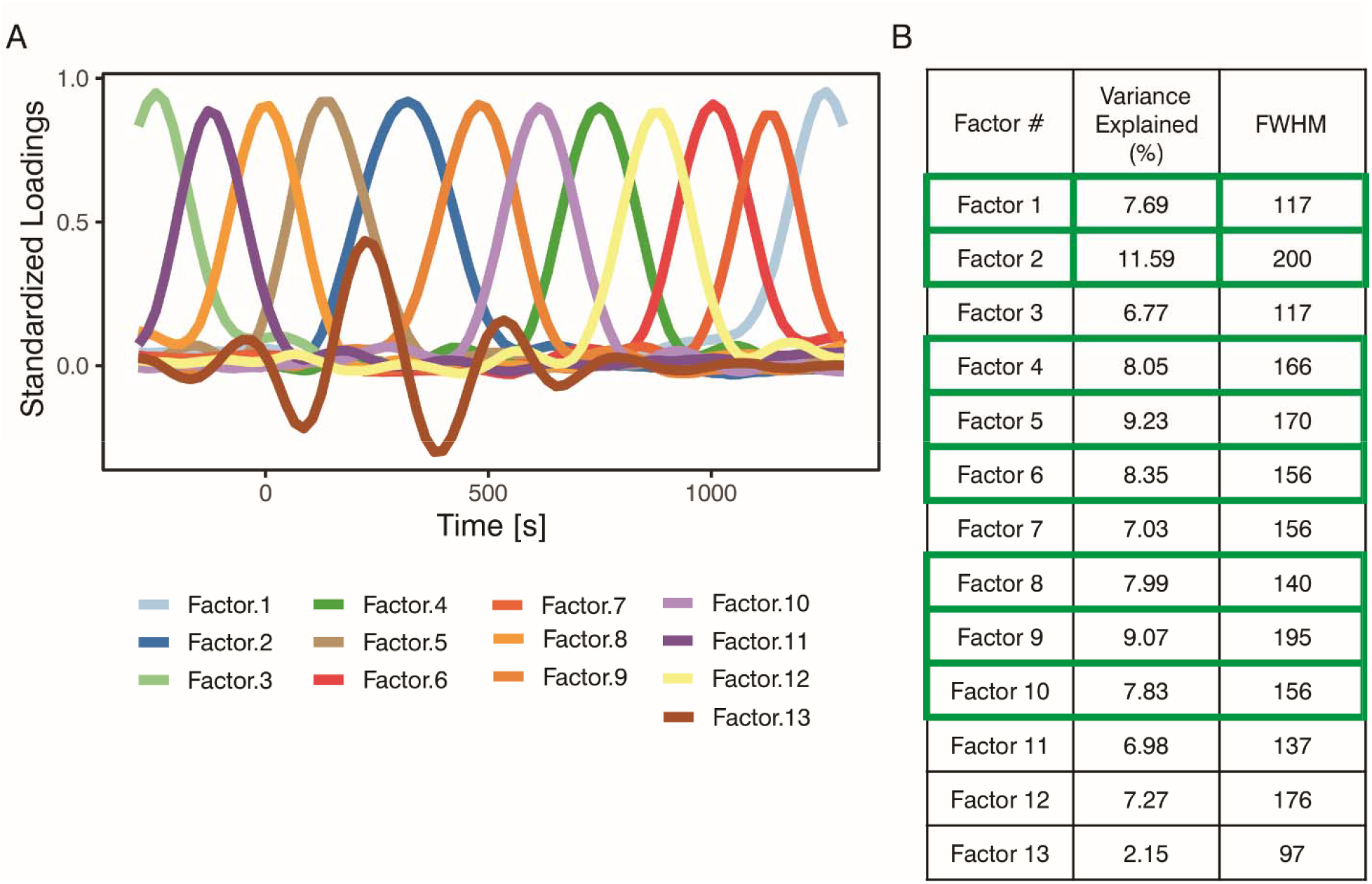
Results of the Temporal Principal Component Analysis. Distribution of time-wise standardized loadings after varimax rotation (Panel A) and percentage of variance explained by each component (Panel B). Components in Panel B are ordered from highest to lowest explained variance. Those outlined in green accounted for more than 1/13 of the variance and were retained for further analysis. FWHM, Full Width at Half Maximum.

#### 3.2.3 Spectral Slope Analysis

Paired *t*-tests were conducted to assess global (condition-average) changes between the pre-stimulus and post-stimulus periods during the trial, while repeated-measures ANOVA was used to examine the CE and CSE. These analyses were performed on the spatial scores for the three spatial components—occipital, frontal, and central (see *Spatial PCA* section)—with their values averaged across nine time windows

(see *Temporal PCA* section). **Figure 6** illustrates the overall time course of aperiodic activity across conditions before and after PCA decomposition, providing an overview of stimulus-induced changes. Because the global post-stimulus negative shift is large, condition-related differences are difficult to discern; therefore, the experimental effects are presented in **Figure 7**. Inspection of the PCA-derived components (**Figure 6B**) reveals apparent offsets between their time courses, with the central and occipital components seemingly shifted upward relative to the frontal component. These offsets likely reflect the way PCA components are constructed—as weighted combinations of signals across all electrodes—rather than activity arising from spatially isolated electrode groups. Accordingly, the labels “frontal,” “central,” and “occipital” refer to the spatial distribution of component loadings, not to distinct neural sources or discrete electrode clusters. The absolute level of each component therefore depends on its specific weighting profile, and differences in offset should not be interpreted as straightforward regional differences in signal magnitude. In the present data, the frontal component appears to capture a relatively larger share of the overall negative slope shift, which may contribute to the apparent upward shift of the remaining components. As expected, the spectral slope exhibited a widespread negativity, with all values remaining below zero. Exploratory paired *t*-test analyses of global (condition-averaged) stimulus-induced changes (**Figure 8** and **Table 1**) showed that, relative to the pre-stimulus period, the stimulus elicited an additional negative shift across the entire post-stimulus interval for the central component, and across all post-stimulus time windows except the final one for the frontal and occipital components. These findings were corroborated by time-resolved cluster-based permutation analyses, which revealed robust post-stimulus deviations from baseline across all spatial components (see Figure 8, orange lines). Specifically, the frontal component exhibited a sustained effect from 8 to 1160 ms (*max*|*Z*| = 5.15, *p_FWER_* < .001), while the occipital component showed a similarly prolonged deviation from 8 to 984 ms (*max*|*Z*| = 6.33, *p_FWER_* < .001). In the central component, two significant clusters were identified: an early modulation between 8 and 145 ms (*max*|*Z*| = 3.58, *p_FWER_*= .003), followed by a long-lasting effect spanning 203 to 1238 ms (*max*|*Z*| = 6.00, *p_FWER_* < .001).

**Figure 6.**
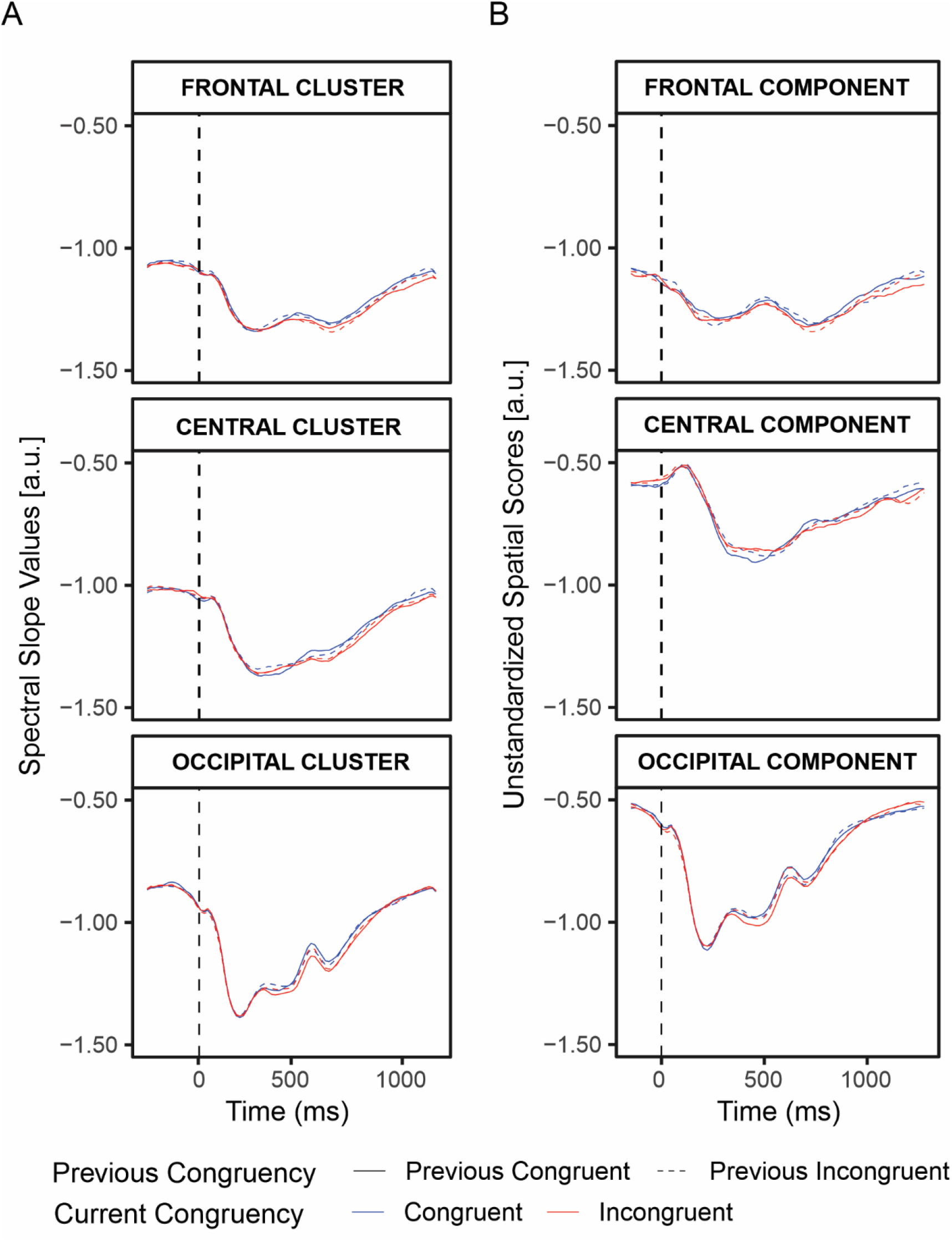
Changes in Aperiodic Activity (Spectral Slope) as a Function of Time. Stimulus-locked spectral slopes for Current Congruency (blue line for congruent, red line for incongruent) by Previous Congruency (solid line for previous congruent, dashed line for previous incongruent), shown for raw data from midline electrodes (Fz, F1 and F2 for frontal cluster; Cz, C1, C2 for central cluster; Oz, O1, O2 for occipital cluster; Panel A) and for key components extracted from spatial PCA (Panel B). The dashed vertical line indicates stimulus onset (time zero). Panel A displays slope values from selected electrodes for illustrative purposes, whereas Panel B presents factor scores derived from all electrodes using PCA weights. Thus, the panels are not directly comparable; rather, they illustrate the relationship between the time course of the raw electrode-level data and the PCA-derived component scores used in the analyses.

**Figure 7.**
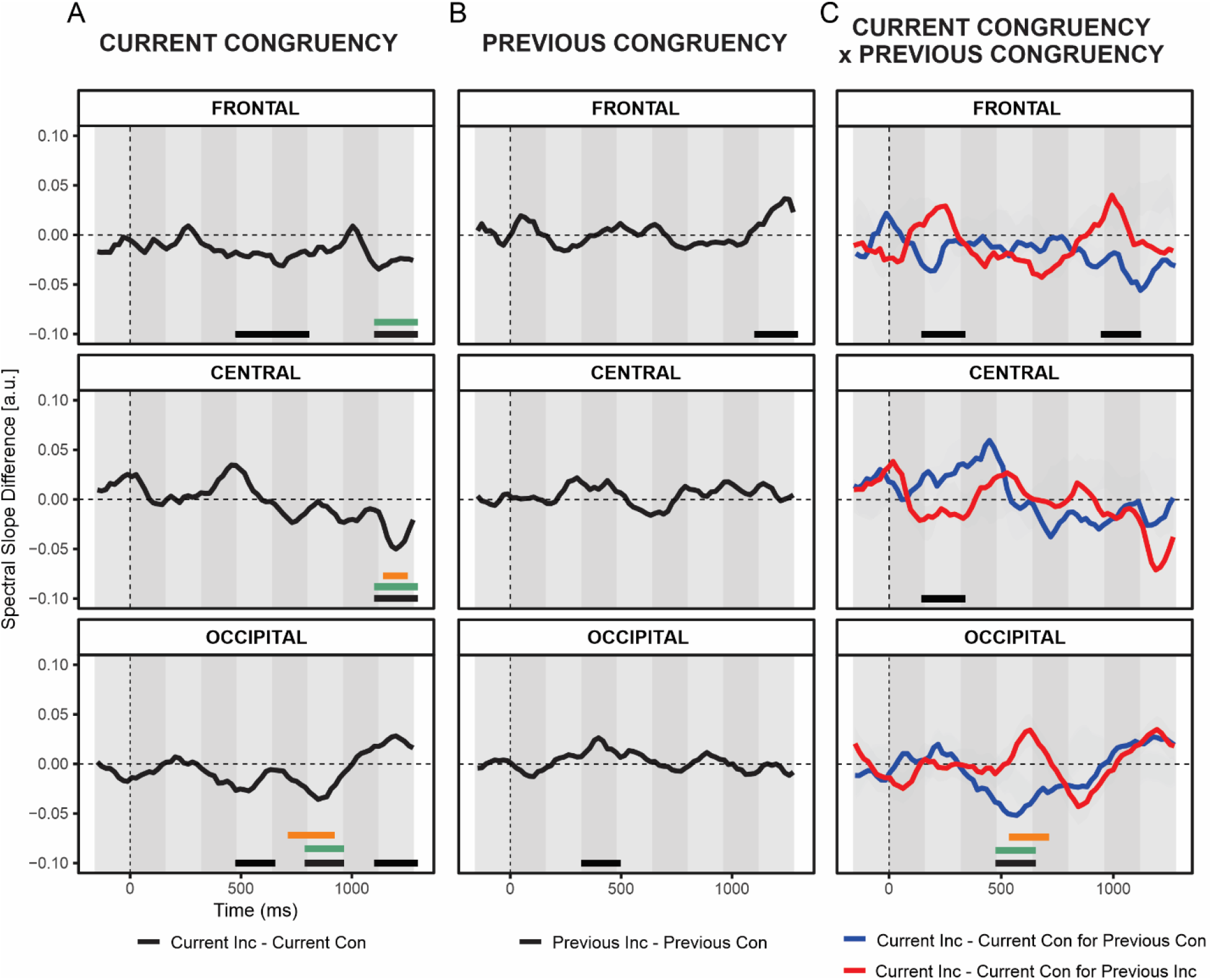
Effects of Experimental Manipulation on the Spectral Slope. The time course of differences in spatial scores for Current Congruency (Panel A), Previous Congruency (Panel B), and their interaction (Panel C) for the frontal (top), central (middle), and occipital (bottom) components derived from spatial PCA. The dashed vertical line indicates stimulus onset (time zero). Shaded areas represent the temporal windows identified by the temporal PCA. Horizontal lines at the bottom of each subplot denote statistically significant effects from the repeated-measures ANOVA (values within the time windows were averaged for testing; no correction for multiple comparisons). Black lines indicate *p* < .05 and green lines indicate *p* < .01 for the ANOVA, whereas orange lines indicate significant clusters identified by the cluster-based permutation analysis across time (*p_FWER_* < .05).

**Figure 8.**
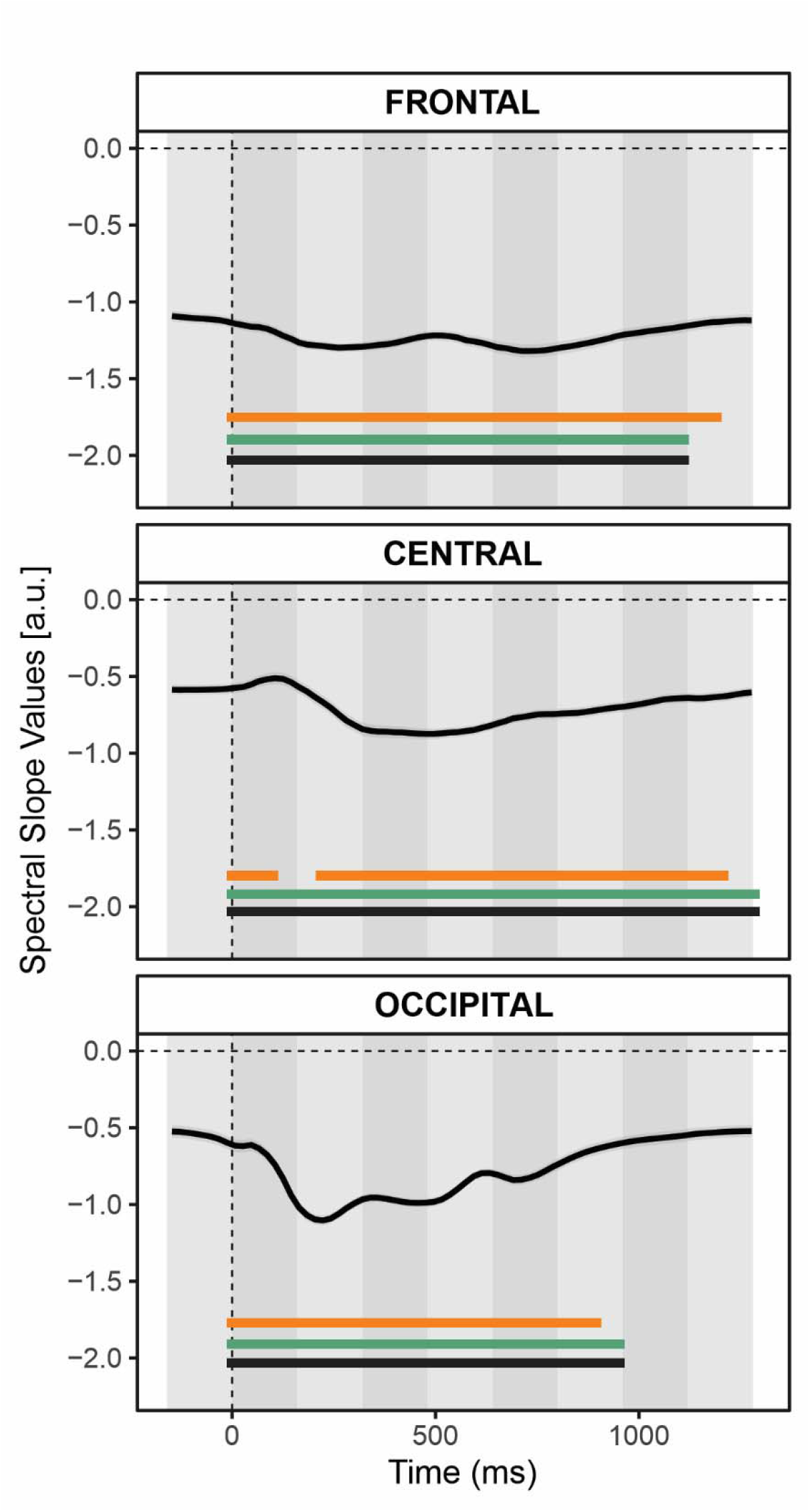
Global Stimulus-Induced Changes in Spectral Slope. The time course of condition-averaged spatial scores for the frontal (top), central (middle), and occipital (bottom) components derived from spatial PCA. The dashed vertical line indicates stimulus onset (time zero). Shaded areas represent the temporal windows identified by the temporal PCA. Horizontal lines at the bottom of each subplot denote statistically significant deviations of post-stimulus values from the pre-stimulus baseline. Black lines indicate *p* < .05; green lines indicate *p* < .01 for the window-based paired *t*-tests (values within the time windows were averaged for testing; no correction for multiple comparisons), and orange lines indicate significant clusters identified by the cluster-based permutation analysis across time (*p_FWER_* < .05).

**Table 1.** Results of t-Tests Assessing Global Changes in Spectral Slope Across Time Windows and Scalp Regions.

| Time<br>Window (ms) | FRONTAL |  |  | CENTRAL |  |  | OCCIPITAL |  |  |
| --- | --- | --- | --- | --- | --- | --- | --- | --- | --- |
|  | *t* | *p* | *d* | *t* | *p* | *d* | *t* | *p* | *d* |
| (0,160] | **-4.23** | **< 0.001** | **-0.60** | **3.67** | **< 0.001** | **0.52** | **-6.88** | **< 0.001** | **-0.98** |
| (160,320] | **-7.44** | **< 0.001** | **-1.06** | **-4.60** | **< 0.001** | **-0.66** | **-14.82** | **< 0.001** | **-2.12** |
| (320,480] | **-6.99** | **< 0.001** | **-1.00** | **-9.08** | **< 0.001** | **-1.30** | **-15.23** | **< 0.001** | **-2.18** |
| (480,640] | **-6.38** | **< 0.001** | **-0.91** | **-10.80** | **< 0.001** | **-1.54** | **-11.84** | **< 0.001** | **-1.69** |
| (640,800] | **-7.65** | **< 0.001** | **-1.09** | **-8.61** | **< 0.001** | **-1.23** | **-9.57** | **< 0.001** | **-1.37** |
| (800,960] | **-7.15** | **< 0.001** | **-1.02** | **-7.06** | **< 0.001** | **-1.01** | **-4.58** | **< 0.001** | **-0.65** |
| (960,1120] | **-4.48** | **< 0.001** | **-0.64** | **-4.88** | **< 0.001** | **-0.70** | -1.13 | 0.262 | -0.162 |
| (1120,1280] | -1.58 | 0.121 | -0.225 | **-2.97** | **0.005** | **-0.42** | 1.07 | 0.288 | 0.154 |

Note. Results of paired *t*-tests comparing post-stimulus values to the pre-stimulus interval (i.e., [-160, 0] ms). For time windows, square brackets indicate inclusion of the boundary value, while parentheses indicate exclusion. Degrees of freedom for all tests are (48). Significant effects ( $p < .05$ , uncorrected for multiple comparisons) are bolded. *d* represents Cohen's *d* effect size.

A series of exploratory repeated-measures ANOVAs assessing the effects of experimental manipulation revealed significant main effects of Current Congruency and Previous Congruency, as well as significant interactions between these two factors, in various time windows and locations (**Figure 7** and **Table 2**).

**Table 2:**
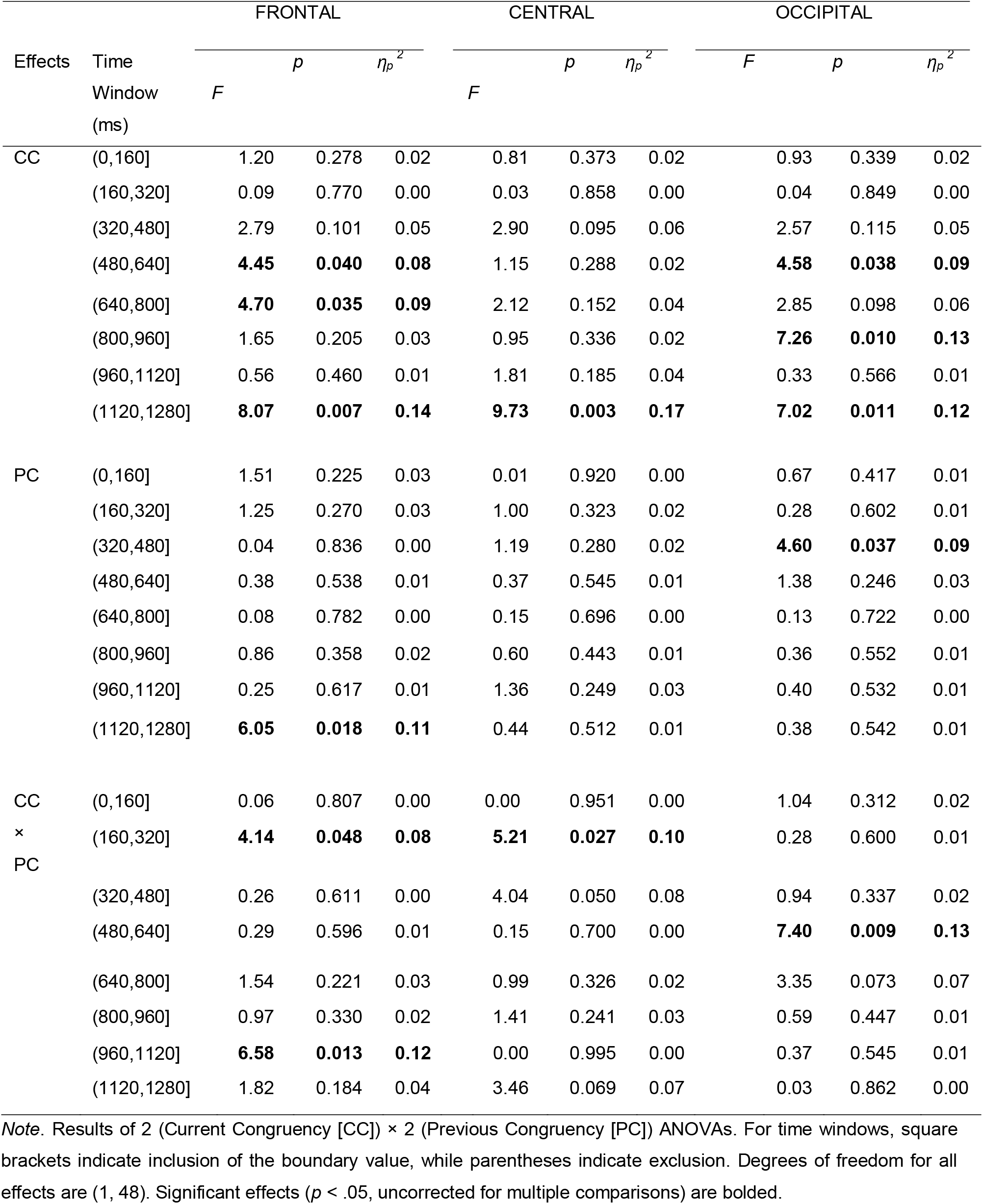
Results of Repeated-Measures ANOVA Assessing the Effects of Experimental Manipulation on Spectral Slope across Time Windows and Scalp Regions.

Significant effects of Current Congruency were observed in three time windows of the frontal component (480–640 ms, 640–800 ms, and 1120–1280 ms), one time window of the central component (1120–1280 ms), and three time windows of the occipital component (480–640 ms, 800–960 ms, and 1120–1280 ms) (see **Figure 7A**). Overall, slope values were more negative for the incongruent condition than for the congruent condition. However, in the final time window of the occipital component (1120–1280 ms), this pattern reversed, with more negative values observed in the congruent than the incongruent condition. Only two significant effects of Previous Congruency were observed (see **Figure 7B**). Slope values were significantly more negative for the previous congruent condition than the previous incongruent condition in the 320—480 ms window for the occipital component and the 1120—1280 ms window for the frontal component. No other statistically significant effects of Previous Congruency were found.

Significant interactions between Current Congruency and Previous Congruency were observed in several time windows across the three components (see **Figure 7C** and **Figure 9**; for the distribution of effects, see **Figure S1**). When considering how previous congruency influenced the current trial, we observed four statistically significant effects in total—three for the current congruent condition and one for the current incongruent condition. In the frontal (160–320 ms, 960–1120 ms) and central (160–320 ms) components, Previous Congruency influenced the current congruent trials. Specifically, in the frontal component, slope values for congruent trials were more negative when preceded by an incongruent trial compared to when preceded by a congruent trial, t_frontal:160–320ms_(48) = 2.33, *p* < .05, *d* = .34, and *t*_frontal:960–1120ms_(48) = 2.17, *p* < .05, *d* = .31. In contrast, the central component showed the opposite pattern: slope values for congruent trials were more negative when preceded by another congruent trial than by an incongruent one, *t*_central:160–320ms_(48) = 2.28, *p* < .05, *d* = .33. No significant effects of Previous Congruency on incongruent trials were found in the frontal or central components (*p*s > .05). However, in the occipital component (480–640 ms), Previous Congruency significantly influenced the current incongruent condition, *t*_occipital:480–640ms_(48) = 3.13, *p* < .05, *d* = .45, with more negative slope values when the incongruent trial was preceded by a congruent one compared to an incongruent one. No effects were observed for the current congruent trials in the occipital component (*p*s > .05). When analyzing the difference between current incongruent and congruent trials separately for each type of preceding trial (i.e., preceded by congruent and preceded by incongruent), we found two significant effects. Specifically, when preceded by a congruent trial, the slope was more negative for current incongruent than for current congruent trials in the late frontal time window (960–1120 ms) and in the occipital window (480–640 ms), *t*_frontal:960–1120ms_(48) = 2.18, *p* < .05, *d* = .32, and *t*_occipital:480–640ms_(48) = 3.57, *p* < .001, *d* = .52. No other significant effects were observed (*p’*s > .05).

**Figure 9.**
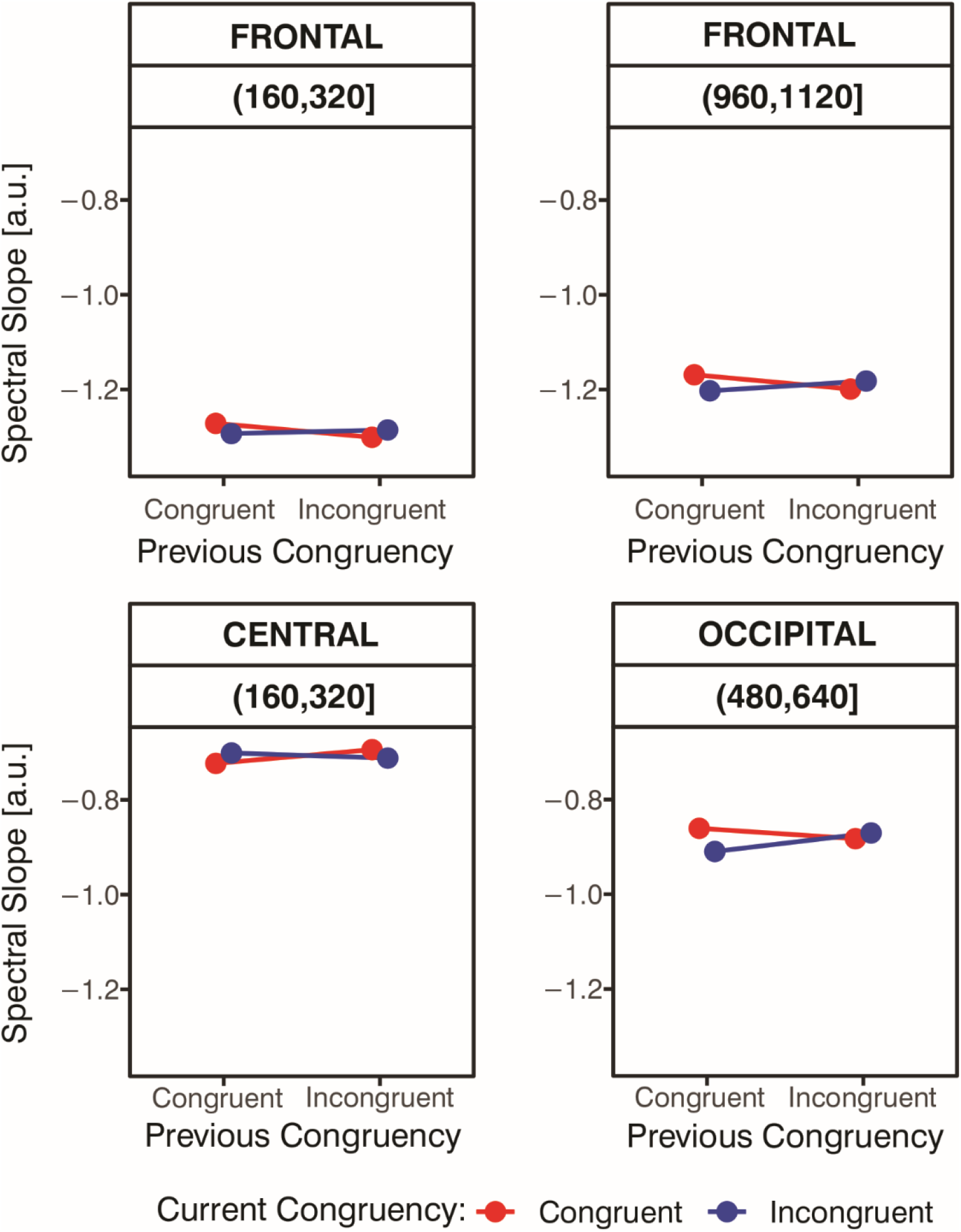
Current Congruency as a Function of Previous Congruency for Spectral Slope. Mean spectral slope values for Current Congruency as a function of Previous Congruency in the time windows and spatial components where significant interactions between these factors were observed: 160–320 ms and 960–1120 ms in the frontal component (top left and top right panels, respectively), 160–320 ms in the central component (bottom left panel), and 480–640 ms in the occipital component (bottom right panel). These outcomes are based on repeated-measures ANOVA conducted within temporal PCA windows and are reported as exploratory analyses without correction for multiple comparisons.

Cluster-based permutation analyses confirmed the robustness of two main effects of Congruency and one interaction effect (see **Figure 7**, orange lines). Specifically, a late main effect of Congruency was observed over the central component (1141–1258 ms; *max*|*Z*| = 3.44, *p_FWER_*= .013). Over the occipital component, a significant main effect of Congruency was found between 770 and 926 ms (*max*|*Z*| = 3.03, *p_FWER_*= .036), as well as a Congruency × Previous Congruency interaction spanning 516–711 ms (*max*|*Z*| = 3.12, *p_FWER_* = .031). No other effects survived FWER correction. Importantly, however, the time-resolved cluster-based permutation results were consistent with the window-based ANOVA findings, identifying effects in comparable temporal ranges (see Table 2). This convergence supports the temporal segmentation derived from the temporal PCA. The full distribution of the *Z*-statistic across time, for both global (condition-averaged) analyses and experimental effects, is presented in the Supplementary Materials (see **Figure S2**).

## 4. Discussion

This study investigated whether and how the aperiodic component of EEG activity—specifically the within-trial time-resolved spectral slope—reflects the CE and CSE during a PWI task. Our findings offer some novel insights into the temporal dynamics and neural mechanisms of cognitive control, demonstrating that fluctuations in post-stimulus aperiodic activity, potentially reflecting shifts in the E/I balance of underlying neural circuits, are linked to the engagement and adaptive modulation of cognitive control.

At the behavioral level, we observed the classic CE, characterized by slower and less accurate responses on incongruent trials compared to congruent ones. We also observed the CSE (Gratton et al., 1992). Specifically, responses to incongruent trials were slower and less accurate when they followed congruent rather than incongruent trials, and likewise, responses to congruent trials were slower and less accurate when they followed incongruent rather than congruent trials. Notably, the CE was statistically significant only following congruent trials but not following incongruent ones, consistent with the idea that experiencing conflict triggers temporary adjustments in control that reduce susceptibility to interference on subsequent trials (Botvinick et al., 2001; Gratton et al., 1992; Grant & Weissman, 2019; 2023; Weissman, 2019).

At the EEG level, we quantified the aperiodic spectral slope in a time-resolved manner using a wavelet approach to capture the dynamics of neural processing. This high-temporal-resolution method allowed us to track rapid fluctuations in aperiodic activity that traditional time-averaged analyses might overlook. The resulting single-cycle estimates should not, however, be interpreted as truly instantaneous slope measures because their temporal support differs across frequencies. Nevertheless, a supplementary simulation indicated that this approach was more sensitive than fixed 500- or 1000-ms FFT windows to brief imposed changes resembling the durations of the FWER-corrected empirical effects, although none of the estimators provided an unbiased instantaneous reconstruction of the imposed slope profile (see **Figure S3**). To reduce data dimensionality, we then applied two PCAs: a spatial PCA and a temporal PCA. The spatial PCA yielded three interpretable components corresponding to occipital, frontal, and central topographies. The temporal PCA provided a resolution of ∼160 ms, segmenting each epoch into eight non-overlapping post-stimulus time windows. As such, subsequent exploratory ANOVA and *t*-test analyses focused on the three spatial components (i.e., frontal, central and occipital) across these windows. These exploratory results were further supplemented by time-resolved cluster-based permutation analyses to assess the robustness of the observed effects.

Visual inspection of the spectral slope time course revealed a widespread negative deflection, with slope values consistently remaining below zero across the scalp (**Figure 6**). This is in line with the expectation that the power spectrum of neural data is negative-going under normal physiological circumstances (Brake et al., 2024; He, 2014; Gao et al., 2017). Subsequent analysis of global (condition-averaged) changes induced by the stimulus showed that the stimulus further increased the negativity of the slope relative to the baseline (pre-stimulus) period (**Figure 8**), in line with findings of post-stimulus steepening of the spectrum by Gyurkovics et al. (2022) and Kałamała et al. (2024). Here, this effect—a negative spectral shift—was particularly robust in the frontal and central components, where it persisted across nearly the entire post-stimulus period, adding some spatial specificity to previous reports. In contrast, the effect in the occipital component was present across most time windows, except the final ones. These findings suggest that the stimulus induced a shift in the aperiodic background activity, presumably reflecting a sustained state of increased cortical inhibition during task engagement. Such an interpretation could align with the well-established pattern of Default Mode Network (DMN) deactivation observed during externally oriented, goal-directed tasks (<u>Buckner et al., 2008</u>; Shulman et al.1997; Mazoyer et al. 2001). The DMN typically shows reduced activity when attention is directed toward goal-oriented behavior, reflecting the suppression of internally directed cognition to support efficient cognitive control and performance (Raichle et al., 2015). Thus, the observed increase in cortical inhibition may represent a neurophysiological correlate of this large-scale network reconfiguration that accompanies engagement in externally focused cognitive demands.

The spectral slope was further modulated by current congruency (**Figure 7A**). In line with our hypothesis, steeper (more negative) slopes were observed during incongruent trials compared to congruent ones in several time windows. The exploratory analysis revealed statistically significant effects in the frontal time intervals of 480–640 ms, 640–800 ms, and 1120–1280 ms after stimulus onset. A similar effect was observed in one relatively late central time interval (1120–1280 ms), as well as in occipital time intervals from 480 to 640 ms and from 800 to 960 ms. Interestingly, in the final occipital time interval (1120–1280 ms), a different pattern emerged, with steeper slopes observed in the congruent than in the incongruent condition. Importantly, the congruency effects identified in the central component (1120–1280 ms) and in the middle occipital time window (800–960 ms) survived FWER correction, as they corresponded to effects detected in the time-resolved cluster-based permutation analysis (central: 1141–1258 ms; occipital: 770–926 ms).

As described in the Introduction, spectral slopes are thought to index E/I balance within synaptic circuits (Ahmad et al., 2022; Gao et al., 2017; Waschke et al., 2021). The presentation of a stimulus, which engages in a task, can lead to a temporary steepening of the spectral slope. Such changes are interpreted as reflecting the increased recruitment of inhibitory mechanisms, which help regulate excessive neural activity and thereby support more efficient cognitive performance (Gratton, 2018). In line with this interpretation, the steeper slopes observed during incongruent compared to congruent trials may reflect a conflict-related neural response that filters out irrelevant signals to support efficient response selection. This may serve as an index of reactive control wherein neural activity is modulated after conflict detection to support accurate performance. Indeed, congruency-related differences in slope were observed relatively late after stimulus onset (around 500 ms), suggesting that control processes reflected in the aperiodic background were engaged *after* conflict detection. The increased cortical inhibition (and/or reduction in cortical excitability) could serve as a neural filtering mechanism, suppressing interference from task-irrelevant signals and enhancing the precision of task-relevant processing, as observed by Gyurkovics et al. (2022) and Kałamała et al. (2024). This mechanism would support more efficient conflict resolution and facilitate more accurate response selection. The proposed interpretation aligns with theoretical accounts positing that an increase in spectral steepness during incongruent trials may indicate a shift toward slower activity, which has been linked to greater engagement of control mechanisms (Gratton, 2018).

Notably, in the final occipital time interval (1120–1280 ms), slopes were less negative for incongruent than for congruent trials, opposite to the expected pattern. Although this finding does not align with our hypothesis, one possible interpretation is that it reflects differences in the functional roles of cortical regions. Occipital areas are primarily involved in sensory processing and feature analysis, which may be relatively more pronounced during congruent trials, whereas frontal and central regions are more strongly engaged in top-down control and conflict resolution, which are typically heightened during incongruent trials. The presence of relatively steeper slopes for congruent trials in this late occipital window may therefore suggest that sensory processing either re-emerges after conflict resolution or becomes more detectable once control-related demands subside. At the same time, a more parsimonious explanation is that this effect reflects a false positive, as it was observed only in the exploratory analyses and was not confirmed by the cluster-based permutation results. It should therefore be interpreted with caution. Taken together, these findings tentatively point to a dynamic interplay between sensory-driven and control-related processes, although future work is needed to replicate this effect and clarify its underlying mechanisms.

The exploratory analysis showed dynamic, trial-to-trial changes in spectral slope steepness across frontal, central, and occipital components, suggesting a modulating effect of the previous congruency on the current congruency (CSE; **Figure 7C**). In the frontal component (160–320 ms, 960–1120 ms), congruent trials following incongruent trials showed more negative slopes than those following congruent trials. In the central component (160–320 ms), the opposite pattern emerged: more negative slopes were observed when congruent trials followed congruent trials. In the occipital component (480–640 ms), incongruent trials preceded by congruent trials again showed more negative slopes than those preceded by another incongruent trial. However, when applying cluster-based permutation analyses to control for multiple comparisons across time, only the occipital effect was replicated (516–711 ms), whereas the frontal and central effects did not survive correction. Thus, the latter findings should be interpreted with caution and primarily considered exploratory. Nevertheless, given that this is the first paper to apply a time-resolved approach to spectral slope data in the adaptive control literature, we provide a tentative interpretation of all observed effects below.

The frontal and occipital patterns were consistent with our predictions: congruent trials preceded by another congruent trial showed the flattest spectral slopes, reflecting minimal engagement of inhibitory processes, whereas incongruent trials following a congruent trial exhibited the steepest slopes, indicating a stronger shift toward inhibition

Furthermore, at these two spatial components, the difference between current congruent and incongruent trials was not observed when trials followed incongruent trials, similarly to our behavioral data where the congruency effect was significant only following the congruent trials, suggesting a CSE pattern. Highlighting the usefulness of a time-resolved approach, these effects emerged at two distinct latencies, with the occipital effect following the earlier frontal effect, indicating a temporal progression of conflict-related processing across cortical regions. Following the E/I balance framework (where steeper slopes indicate greater inhibition/lower excitation) and the CE interpretation proposed above, the observed patterns at frontal and occipital sites jointly suggest that reduced inhibition of synaptic circuits occurred when conflict (i.e., interference from irrelevant information) was minimal, whereas increased inhibition became evident as the level of conflict increased (this interpretation is reported in the model depicted in **Figure 10**).

**Figure 10.**
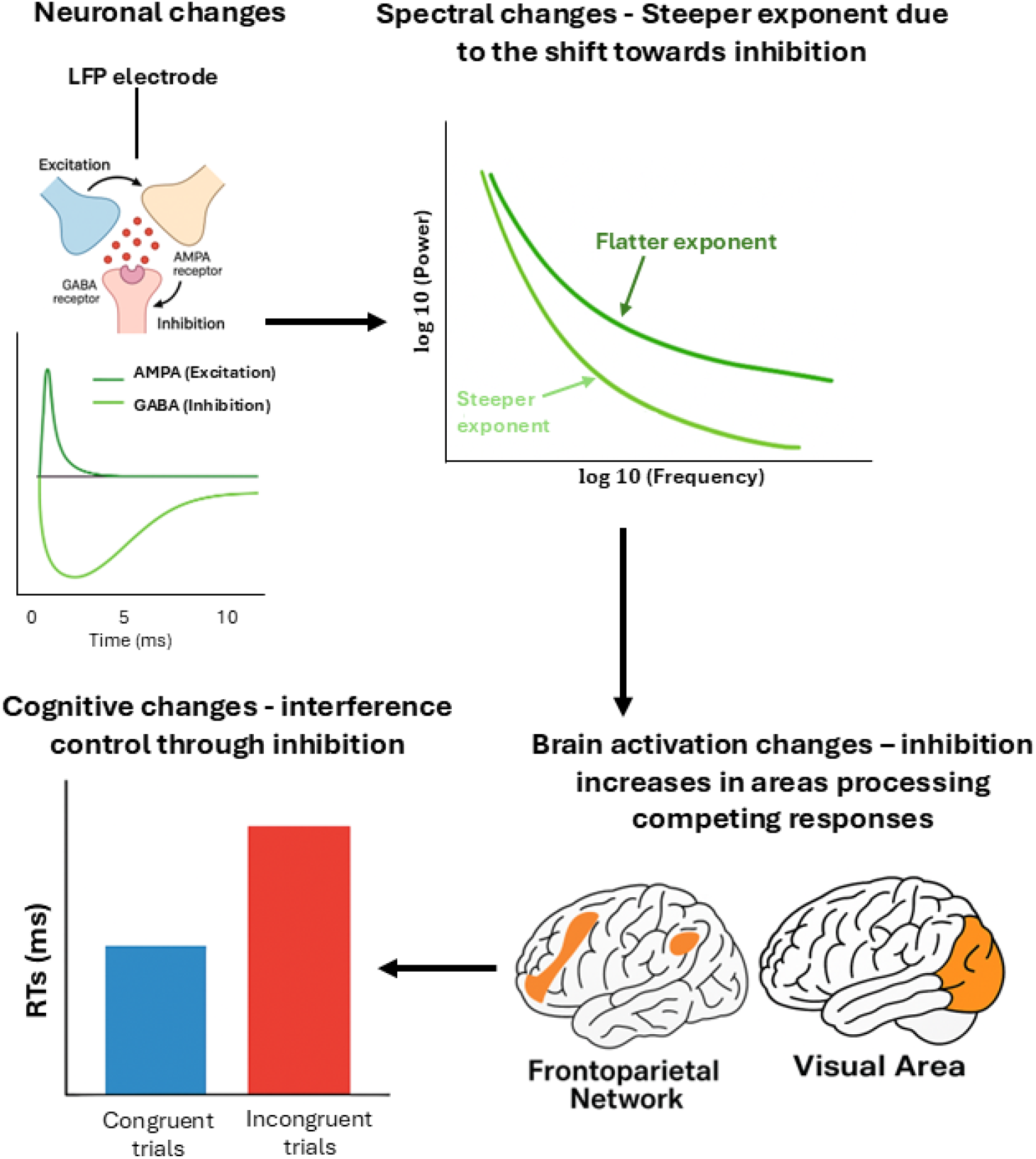
From neuronal inhibition to cognitive control level. Schematic overview of the proposed mechanisms linking neuronal-level changes in excitation and inhibition to cognitive control. (Top left) A shift in the local excitation/inhibition balance, captured by local field potentials (LFPs), reflects increased GABAergic inhibition relative to AMPA-mediated excitation. (Top right) This shift could lead to spectral changes characterized by a steeper 1/*f* slope in the power spectrum. (Bottom right) At the brain-network level, increased inhibition manifests as enhanced activation in regions involved in resolving competing responses. (Bottom left) At the cognitive level, stronger cognitive control supports performance in interference tasks, reflected by increased reaction times (RTs) for incongruent compared to congruent trials.

This interpretation aligns with the conflict monitoring theory (Botvinick et al., 2001) described in the Introduction. In our data, occipital effects occurred later than frontal changes, consistent with a sequence in which conflict is detected in frontal (possibly ACC-related) regions and subsequently resolved through top-down modulation of occipital sensory processing.

However, frontal and central components showed opposing patterns within the same 160–320 ms window. A potential explanation for this inconsistency is that the spectral slope may reflect multiple underlying mechanisms and/or neural substrates, with their relative contributions varying across scalp locations. As discussed below, our design was not entirely free of potential confounds, such as those related to feature integration. Therefore, one possibility is that frontal activity reflects the recruitment of the frontoparietal network during cognitive control processes (Niendam et al., 2012; Gratton, Sun, & Petersen, 2018). In contrast, the central pattern may reflect the engagement of a different network, such as the DMN. From this perspective, frontal activity would primarily signal the implementation of cognitive control, whereas central activity might capture the contribution of additional task-related mechanisms. At the same time, a more parsimonious interpretation is that at least one of these effects may reflect a false positive, as these effects did not survive correction for multiple comparisons in the cluster-based permutation analysis. This raises the possibility that the apparent dissociation between frontal and central components may be driven, at least in part, by noise rather than by distinct underlying neural processes. Given the relatively large number of comparisons in the exploratory analyses, such spurious effects are statistically plausible. Accordingly, these findings should be interpreted with caution and treated as hypothesis-generating rather than confirmatory. Future studies using higher-powered designs and independent replication will be necessary to determine whether these patterns reflect reliable and functionally meaningful differences across cortical regions.

## 5. Limitations and Future Directions

This study is the first to examine the time-resolved aperiodic EEG in relation to cognitive control. While it provides novel insights, it also has certain limitations that point to directions for future research. First, the PWI task does not involve a fully confound-minimized design, making it difficult to disentangle feature integration effects from cognitive control engagement. This limitation may confound the interpretation of the CSE or increase the variability in the data, as feature integration processes can, at least to some extent, obscure genuine adjustments in cognitive control (e.g., Duthoo et al., 2014; Hommel et al., 2004; Mayr et al., 2003). Future research should adopt confound-minimized designs that disentangle feature integration from more traditional cognitive control processes to clarify the underlying mechanisms (Braem et al., 2019; Schmidt & Liefooghe, 2016; Weissman et al., 2014). It should be noted, however, that feature integration and cognitive control likely operate in concert rather than in opposition to one another (e.g., Abrahamse et al., 2016).

Second, the observed aperiodic effects were relatively small (partial η*_p_²* = 0.08–0.14), with three effects surviving correction for multiple comparisons (FWER). The exploratory findings should therefore be interpreted with caution. Several factors may account for the limited magnitude of the effects: (a) the spectral slope reflects a relatively stable property of neural activity, which typically fluctuates within a constrained range, making large condition-related shifts inherently unlikely; (b) the time-resolved approach prioritizes temporal precision, but this comes at the cost of reduced signal-to-noise compared to analyses that aggregate data over broader time intervals; (c) the experimental task involved multiple concurrent manipulations; in addition to Congruency and Previous Congruency, the task included factors such as Stimulus Repetition, Category Repetition, and Response–Stimulus Interval, which, although not the focus of the present analyses, may nonetheless have influenced participants’ processing. As a result, the observed aperiodic dynamics likely reflect the combined contribution of several overlapping processes, thereby attenuating the apparent effect of any single manipulation.

It is also possible that the sample size was not sufficient to reliably detect these effects. While our sample size (*N* = 49) is typical for EEG research, future studies with larger samples are needed to replicate these findings and establish their robustness and generalizability (Clayson et al., 2019). Moreover, as this is one of the first studies to apply a time-resolved approach to the aperiodic data, there was no prior basis for determining the optimal sample size for detecting these effects, underscoring the need for follow-up work. Finally, the present findings were obtained using the Picture–Word Interference (PWI) task, which primarily engages lexical–semantic processing with concrete stimuli. It remains an open question whether similar time-resolved aperiodic dynamics would be observed in other cognitive control paradigms, such as task switching, flanker, or Stroop tasks (von Bastian et al., 2020), which typically involve more abstract stimuli and place different demands on control processes. Moreover, it should be acknowledged that the spectral slope could also reflect fluctuations in arousal (Mocchi et al., 2024), or motor preparation (Wilson et al., 2022), particularly given the temporal proximity of the observed CE effects to response-related activity. The spectral slope may also reflect different processes—or combinations of processes—depending on the cortical region analyzed, underscoring a broader limitation of surface-level aperiodic EEG studies. We speculated about distinct functional interpretations of the spectral slope across scalp distributions. This reasoning parallels long-standing findings in ERP research, where different components arise from distinct neural generators with specific spatiotemporal and functional properties (Fabiani et al., 2007). To disentangle these contributions, future studies could combine aperiodic analyses with source localization techniques or with techniques with better spatial resolution (e.g., EEG–fMRI) to better identify the cortical generators and functional significance of slope changes. Additionally, experimental manipulations targeting specific mechanisms—such as pharmacological modulation of the E/I balance or controlled changes in arousal—could help clarify the functional specificity of aperiodic dynamics.

## 6. Conclusions

The time-resolved analysis of the aperiodic EEG activity presented here offers a novel approach to studying cognitive control, showing how inhibitory dynamics evolve throughout the trial. This fine-grained approach demonstrates that aperiodic activity can index subtle changes in conflict resolution and can capture adjustments in cognitive control processes. Specifically, our results show slope modulations during incongruent trials consistent with increased inhibitory activity relative to congruent trials, which may be linked to control mechanisms, such as the suppression of irrelevant representations. In parallel, adjustments in cognitive control driven by the previous trial’s context were reflected in slope modulations, emerging most robustly in the occipital component, with weaker and less consistent effects observed in frontal and central regions. The sequence of effects across spatial components was broadly consistent with the temporal unfolding of the CSE (Botvinick et al., 2001). These CSE-related effects suggest that the spectral slope is sensitive not only to the immediate demands of the task, but also to internal control states carried over from preceding trials, highlighting its potential as a marker of proactive adjustments in cortical excitability. Overall, this time-resolved framework highlights that the aperiodic component could capture the spatial-temporal dynamics of cognitive control, offering a novel account of how the brain resolves conflict and flexibly adapts control in real time.

## Acknowledgmentsx

This work was partly supported by NIA grant RF1AG062666 to M. Fabiani and G. Gratton. This work was conducted in partial fulfillment of the Ph.D. degree requirements of Virginia Tronelli.

## Data Availability Statement

Data and code necessary to reproduce the statistical analyses (section 3.2.3) will be available at https://osf.io/z3qsg.

*Link for reviewers*: https://osf.io/z3qsg/overview?view_only=fc282671ba704386bf47f7776d4e4509

## Ethics approval statement

The experimental protocol adhered to the principles outlined in the Declaration of Helsinki and received approval from the Ethical Committee of the University of Bologna. Written informed consent was obtained from all participants.

## Author Contributions

Conceptualization: Virginia Tronelli, Patrycja Kałamała, Gabriele Gratton, Monica Fabiani; Maurizio Codispoti, Andrea De Cesarei

Methodology: Patrycja Kałamała, Gabriele Gratton, Mate Gyurkovics

Software: Patrycja Kałamała, Mate Gyurkovics

Data curation: Virginia Tronelli, Kathy A. Low

Investigation: Virginia Tronelli

Formal analysis: Patrycja Kałamała, Virginia Tronelli

Supervision: Gabriele Gratton, Monica Fabiani, Maurizio Codispoti, Andrea De Cesarei

Funding acquisition: Gabriele Gratton, Monica Fabiani

Visualization: Patrycja Kałamała, Virginia Tronelli

Project administration: Monica Fabiani, Kathy A. Low

Resources: Gabriele Gratton, Monica Fabiani, Kathy A. Low, Maurizio Codispoti, Andrea De Cesarei

Writing - original draft: Patrycja Kałamała, Virginia Tronelli

Writing - review & editing: Virginia Tronelli, Patrycja Kałamała, Gabriele Gratton, Monica Fabiani, Mate Gyurkovics, Kathy A. Low, Maurizio Codispoti, Andrea De Cesarei

## Footnotes

1 We thank the Reviewers for suggesting the use of cluster-based permutation analysis, which strengthened the statistical rigor of the study.

## References

Abrahamse, E. L., Braem, S., Notebaert, W., & Verguts, T. (2016). Grounding cognitive control in associative learning. Psychological Bulletin, 142(7), 693–728. 10.1037/bul0000047

Akbarian, F., Rossi, C., Costers, L., D’hooghe, M. B., D’haeseleer, M., Nagels, G., & Van Schependom, J. (2024). Stimulus-related modulation in the 1/f spectral slope suggests impaired inhibition during a working memory task in people with multiple sclerosis. Multiple Sclerosis Journal, 30(8), 1036–1046. 10.1177/13524585241253777

Ahmad, J., Ellis, C., Leech, R., Voytek, B., Garcés, P., Jones, E., et al. (2022). From mechanisms to markers: Novel noninvasive EEG proxy markers of the neural excitation and inhibition system in humans. Translational Psychiatry, 12(1), 467. 10.1038/s41398-022-02218-z

Botvinick, M. M., Braver, T. S., Barch, D. M., Carter, C. S., & Cohen, J. D. (2001). Conflict monitoring and cognitive control. Psychological Review, 108(3), 624–652. 10.1037/0033-295X.108.3.624

Boudewyn, M. A., Erickson, M. A., Winsler, K., Barch, D. M., Carter, C. S., Frank, M. J., et al. (2025). Assessing trial-by-trial electrophysiological and behavioral markers of attentional control and sensory precision in psychotic and mood disorders. Schizophrenia Bulletin, 51(2), 543–555. 10.1093/schbul/sbae038

Brake, N., Duc, F., Rokos, A., Arseneau, F., Shahiri, S., Khadra, A., & Plourde, G. (2024). A neurophysiological basis for aperiodic EEG and the background spectral trend. Nature Communications, 15(1). 10.1038/s41467-024-45922-8

Braeken, J., & Van Assen, M. A. (2017). An empirical Kaiser criterion. Psychological Methods, 22(3), 450–466. 10.1037/met0000074

Braem, S., Bugg, J. M., Schmidt, J. R., Crump, M. J. C., Weissman, D. H., Notebaert, W., & Egner, T. (2019). Measuring adaptive control in conflict tasks. Trends in Cognitive Sciences, 23(9), 769–783. 10.1016/j.tics.2019.07.002

Buckner, R. L., Andrews-Hanna, J. R., & Schacter, D. L. (2008). The brain’s default network: Anatomy, function, and relevance to disease. Annals of the New York Academy of Sciences, 1124(1), 1–38. 10.1196/annals.1440.011

Cavanagh, J. F., & Frank, M. J. (2014). Frontal theta as a mechanism for cognitive control. Trends in Cognitive Sciences, 18(8), 414–421. 10.1016/j.tics.2014.04.012

Chini, M., Pfeffer, T., & Hanganu-Opatz, I. (2022). An increase of inhibition drives the developmental decorrelation of neural activity. eLife, 11, e78811. 10.7554/eLife.78811

Clayson, P. E., Carbine, K. A., Baldwin, S. A., & Larson, M. J. (2019). Methodological reporting behavior, sample sizes, and statistical power in studies of event-related potentials: Barriers to reproducibility and replicability. Psychophysiology, 56(11), e13437. 10.1111/psyp.13437

Cohen, M. X., & Donner, T. H. (2013). Midfrontal conflict-related theta-band power reflects neural oscillations that predict behavior. Journal of Neurophysiology, 110(12), 2752–2763. 10.1152/jn.00479.2013

Cousineau, D. (2005). Confidence intervals in within-subject designs: A simpler solution to Loftus and Masson’s method. Tutorials in Quantitative Methods for Psychology, 1(1), 42–45. 10.20982/tqmp.01.1.p042

De Cesarei, A., Cavicchi, S., Micucci, A., & Codispoti, M. (2019). Categorization goals modulate the use of natural scene statistics. Journal of Cognitive Neuroscience, 31(1), 109–125. 10.1162/jocn_a_01333

De Cesarei, A., Cavicchi, S., Cristadoro, G., & Lippi, M. (2021). Do humans and deep convolutional neural networks use visual information similarly for the categorization of natural scenes? Cognitive Science, 45, e13009. 10.1111/cogs.13009

De Cesarei, A., D’Ascenzo, S., Nicoletti, R., & Codispoti, M. (2023). Novelty and learning in cognitive control: Evidence from the Simon task. Psychological Research, 87, 2390–2406. 10.1007/s00426-023-01813-z

Delorme, A., & Makeig, S. (2004). EEGLAB: An open-source toolbox for analysis of single-trial EEG dynamics including independent component analysis. Journal of Neuroscience Methods, 134(1), 9–21. 10.1016/j.jneumeth.2003.10.009

Donoghue, T., Haller, M., Peterson, E. J., Varma, P., Sebastian, P., Gao, R., et al. (2020). Parameterizing neural power spectra into periodic and aperiodic components. Nature Neuroscience, 23(12), 1655–1665. 10.1038/s41593-020-00744-x

Duthoo, W., Abrahamse, E. L., Braem, S., Boehler, C. N., & Notebaert, W. (2014). The heterogeneous world of congruency sequence effects: An update. Frontiers in Psychology, 5, 1001. 10.3389/fpsyg.2014.01001

Egner, T. (2007). Congruency sequence effects and cognitive control. Cognitive, Affective, & Behavioral Neuroscience, 7(4), 380–390. 10.3758/CABN.7.4.380

Egner, T. (2023). Principles of cognitive control over task focus and task switching. Nature Reviews Psychology, 2(11), 702–714. 10.1038/s44159-023-00234-4

Eriksen, B. A., & Eriksen, C. W. (1974). Effects of noise letters upon the identification of a target letter in a nonsearch task. Perception & Psychophysics, 16(1), 143–149. 10.3758/BF03203267

Fabiani, M., Gratton, G., & Federmeier, K. D. (2007). Event-related brain potentials: Methods, theory, and applications. In Handbook of psychophysiology (3rd ed., pp. 85–119). Cambridge University Press. 10.1017/CBO9780511546396.004

Frelih, T., Matkovič, A., Mlinarič, T., Bon, J., & Repovš, G. (2025). Modulation of aperiodic EEG activity provides sensitive index of cognitive state changes during working memory task. eLife, 13. 10.7554/eLife.101071.3

Gao, R., Peterson, E. J., & Voytek, B. (2017). Inferring synaptic excitation/inhibition balance from field potentials. NeuroImage, 158, 70–78. 10.1016/j.neuroimage.2017.06.078

Grant, L. D., & Weissman, D. H. (2019). Turning distractors into targets increases the congruency sequence effect. Acta Psychologica, 192, 31–41. 10.1016/j.actpsy.2018.10.010

Grant, L. D., & Weissman, D. H. (2023). The binary structure of event files generalizes to abstract features: A nonhierarchical explanation of task set boundaries for the congruency sequence effect. Journal of Experimental Psychology: Learning, Memory, and Cognition, 49(7), 1033–1050. 10.1037/xlm0001148

Gratton, C., Sun, H., & Petersen, S. E. (2018). Control networks and hubs. Psychophysiology, 55(3), e13032. 10.1111/psyp.13032

Gratton, G. (2018). Brain reflections: A circuit-based framework for understanding information processing and cognitive control. Psychophysiology, 55(3), e13038. 10.1111/psyp.13038

Gratton, G., Coles, M. G. H., & Donchin, E. (1983). A new method for off-line removal of ocular artifact. Electroencephalography and Clinical Neurophysiology, 55(4), 468–484. 10.1016/0013-4694(83)90135-9

Gratton, G., Coles, M. G. H., & Donchin, E. (1992). Optimizing the use of information: Strategic control of activation of responses. Journal of Experimental Psychology: General, 121(4), 480–506. 10.1037/0096-3445.121.4.480

Gratton, G., Cooper, P., Fabiani, M., Carter, C. S., & Karayanidis, F. (2018). Dynamics of cognitive control: Theoretical bases, paradigms, and a view for the future. Psychophysiology, 55(3), e13016. 10.1111/psyp.13016

Gyurkovics, M., Clements, G. M., Low, K. A., Fabiani, M., & Gratton, G. (2022). Stimulus-induced changes in 1/f-like background activity in EEG. Journal of Neuroscience, 42(37), 7144–7151. 10.1523/JNEUROSCI.0414-22.2022

Gyurkovics, M., Kovacs, M., Jaquiery, M., Palfi, B., Dechterenko, F., & Aczel, B. (2020). Registered replication report of Weissman, D. H., Jiang, J., and Egner, T. (2014): Determinants of congruency sequence effects without learning and memory confounds. Attention, Perception, & Psychophysics, 82(8), 3777–3787. 10.3758/s13414-020-02021-2

Gyurkovics, M., & Levita, L. (2021). Dynamic adjustments of midfrontal control signals in adults and adolescents. Cerebral Cortex, 31(2), 795–808. 10.1093/cercor/bhaa258

Gyurkovics, M., Stafford, T., & Levita, L. (2020). Cognitive control across adolescence: Dynamic adjustments and mind-wandering. Journal of Experimental Psychology: General, 149(6), 1017–1031. 10.1037/xge0000698

He, B. J. (2014). Scale-free brain activity: Past, present, and future. Trends in Cognitive Sciences, 18(9), 480–487. 10.1016/j.tics.2014.04.003

He, W., Donoghue, T., Sowman, P. F., Seymour, R. A., Brock, J., Crain, S., et al. (2019). Co-increasing neuronal noise and beta power in the developing brain. bioRxiv. 10.1101/839258

Hommel, B. (2004). Event files: Feature binding in and across perception and action. Trends in Cognitive Sciences, 8(11), 494–500. 10.1016/j.tics.2004.08.007

Jasper, H. H. (1958). The ten–twenty electrode system of the International Federation. Electroencephalography and Clinical Neurophysiology, 10, 371–375.

Jia, S., Liu, D., Song, W., Beste, C., Colzato, L., & Hommel, B. (2024). Tracing conflict-induced cognitive-control adjustments over time using aperiodic EEG activity. Cerebral Cortex, 34(5). 10.1093/cercor/bhae185

Jiménez, L., & Méndez, A. (2014). Even with time, conflict adaptation is not made of expectancies. Frontiers in Psychology, 5, 1042. 10.3389/fpsyg.2014.01042

Kaiser, H. F. (1958). The varimax criterion for analytic rotation in factor analysis. Psychometrika, 23(3), 187–200. 10.1007/BF02289233

Kałamała, P., Clements, G. M., Gyurkovics, M., Chen, T., Low, K. A., Fabiani, M., & Gratton, G. (2026). How to improve the reliability of aperiodic parameter estimates in M/EEG: A method comparison. Psychophysiology, 63(3), e70272. 10.1111/psyp.70272

Kałamała, P., Gyurkovics, M., Bowie, D. C., Clements, G. M., Low, K. A., Dolcos, F., Fabiani, M., & Gratton, G. (2024). Event-induced modulation of aperiodic background EEG: Attention-dependent and age-related shifts in E:I balance, and their consequences for behavior. Imaging Neuroscience, 2, 1–18. 10.1162/imag_a_00054

Kim, S., & Cho, Y. S. (2014). Congruency sequence effect without feature integration and contingency learning. Acta Psychologica, 149, 60–66. 10.1016/j.actpsy.2014.03.004

Lopez-Calderon, J., & Luck, S. J. (2014). ERPLAB: An open-source toolbox for the analysis of event-related potentials. Frontiers in Human Neuroscience, 8, 213. 10.3389/fnhum.2014.00213

Lu, R., Dermody, N., Duncan, J., & Woolgar, A. (2024). Aperiodic and oscillatory systems underpinning human domain-general cognition. Communications Biology, 7(1), 1643. 10.1038/s42003-024-07397-7

Manyukhina, V. O., Prokofyev, A. O., Obukhova, T. S., Stroganova, T. A., & Orekhova, E. V. (2024). Changes in high-frequency aperiodic 1/f slope and periodic activity reflect post-stimulus functional inhibition in the visual cortex. Imaging Neuroscience, 2, 1–24. 10.1162/imag_a_00146

Mayr, U., Awh, E., & Laurey, P. (2003). Conflict adaptation effects in the absence of executive control. Nature Neuroscience, 6(5), 450–452. 10.1038/nn1051

Mazoyer, B., Zago, L., Mellet, E., Bricogne, S., Etard, O., Houdé, O., et al. (2001). Cortical networks for working memory and executive functions sustain the conscious resting state in humans. Brain Research Bulletin, 54(3), 287–298. 10.1016/S0361-9230(00)00437-8

Mocchi, M., Bartoli, E., Magnotti, J., de Gee, J. W., Metzger, B., Pascuzzi, B., et al. (2024). Aperiodic spectral slope tracks the effects of brain state on saliency responses in the human auditory cortex. Scientific Reports, 14(1), 30751. 10.1038/s41598-024-80911-3

Myrov, V., Siebenhühner, F., Juvonen, J. J., Arnulfo, G., Palva, S., & Palva, J. M. (2024). Rhythmicity of neuronal oscillations delineates their cortical and spectral architecture. Communications Biology, 7(1), 405. 10.1038/s42003-024-06083-y

Niendam, T. A., Laird, A. R., Ray, K. L., Dean, Y. M., Glahn, D. C., & Carter, C. S. (2012). Meta-analytic evidence for a superordinate cognitive control network subserving diverse executive functions. Cognitive, Affective, & Behavioral Neuroscience, 12, 241– 268. 10.3758/s13415-011-0083-5

Raichle, M. E. (2015). The brain’s default mode network. Annual Review of Neuroscience, 38, 433–447. 10.1146/annurev-neuro-071013-014030

Rosinski, R. R. (1977). Picture-word interference is semantically based. Child Development, 48(2), 643–647. https://www.jstor.org/stable/1128667

Scharf, F., Widmann, A., Bonmassar, C., & Wetzel, N. (2022). A tutorial on the use of temporal principal component analysis in developmental ERP research: Opportunities and challenges. Developmental Cognitive Neuroscience, 54, 101072. 10.1016/j.dcn.2022.101072

Schiltenwolf, M., Kiesel, A., Frings, C., & Dignath, D. (2024). Memory for abstract control states does not decay with increasing retrieval delays. Psychological Research, 88(2), 547–561. 10.1007/s00426-023-01870-4

Schmidt, J. R., & Liefooghe, B. (2016). Feature integration and task switching: Diminished switch costs after controlling for stimulus, response, and cue repetitions. PLOS ONE, 11(3), e0151188. 10.1371/journal.pone.0151188

Schmidt, J. R., & Weissman, D. H. (2014). Congruency sequence effects without feature integration or contingency learning confounds. PLOS ONE, 9(7), e102337. 10.1371/journal.pone.0102337

Shulman, G. L., Fiez, J. A., Corbetta, M., Buckner, R. L., Miezin, F. M., Raichle, M. E., & Petersen, S. E. (1997). Common blood flow changes across visual tasks: II. Decreases in cerebral cortex. Journal of Cognitive Neuroscience, 9(5), 648–663. 10.1162/jocn.1997.9.5.648

Steiner, M. D., & Grieder, S. (2020). EFAtools: An R package with fast and flexible implementations of exploratory factor analysis tools. Journal of Open Source Software, 5(53), 2521. 10.21105/joss.02521

Thomson, G. H. (1938). The factorial analysis of human ability. University of London Press.

Thurstone, L. L. (1935). The vectors of mind: Multiple-factor analysis for the isolation of primary traits. University of Chicago Press. 10.1037/10018-007

Tronelli, V., Codispoti, M., & De Cesarei, A. (2026). Cognitive control during scene categorization: The role of identity repetition and timing in congruence sequence effects. Quarterly Journal of Experimental Psychology, 79(1). 10.1177/17470218251335293

van Maanen, L., van Rijn, H., & Borst, J. P. (2009). Stroop and picture-word interference are two sides of the same coin. Psychonomic Bulletin & Review, 16(6), 987–999. 10.3758/PBR.16.6.987

von Bastian, C. C., Blais, C., Brewer, G., Gyurkovics, M., Hedge, C., Kałamała, P., et al. (2020). Advancing the understanding of individual differences in attentional control: Theoretical, methodological, and analytical considerations. PsyArXiv. 10.31234/osf.io/x3b9k

Voytek, B., & Knight, R. T. (2015). Dynamic network communication as a unifying neural basis for cognition, development, aging, and disease. Biological Psychiatry, 77(12), 1089–1097. 10.1016/j.biopsych.2015.04.016

Voytek, B., Kramer, M. A., Case, J., Lepage, K. Q., Tempesta, Z. R., Knight, R. T., & Gazzaley, A. (2015). Age-related changes in 1/f neural electrophysiological noise. Journal of Neuroscience, 35(38), 13257–13265. 10.1523/JNEUROSCI.2332-14.2015

Waschke, L., Donoghue, T., Fiedler, L., Smith, S., Garrett, D. D., Voytek, B., & Obleser, J. (2021). Modality-specific tracking of attention and sensory statistics in the human electrophysiological spectral exponent. eLife, 10, e70068. 10.7554/eLife.70068

Weissman, D. H. (2019). Let your fingers do the walking: Finger force distinguishes competing accounts of the congruency sequence effect. Psychonomic Bulletin & Review, 26(5), 1619–1626. 10.3758/s13423-019-01626-5

Weissman, D. H., Jiang, J., & Egner, T. (2014). Determinants of congruency sequence effects without learning and memory confounds. Journal of Experimental Psychology: Human Perception and Performance, 40(5), 2022–2037. 10.1037/a0037454

Wilson, L. E., da Silva Castanheira, J., & Baillet, S. (2022). Time-resolved parameterization of aperiodic and periodic brain activity. eLife, 11, e77348. 10.7554/eLife.77348

Yan, J., Yu, S., Mückschel, M., Colzato, L., Hommel, B., & Beste, C. (2024). Aperiodic neural activity reflects metacontrol in task-switching. Scientific Reports, 14(1), 24088. 10.1038/s41598-024-74867-7

Zhang, C., Stock, A. K., Mückschel, M., Hommel, B., & Beste, C. (2023). Aperiodic neural activity reflects metacontrol. Cerebral Cortex, 33(12), 7941–7951. 10.1093/cercor/bhad089

