## Supplementary Materials for "Changes in aperiodic (1/*f* slope) activity during a picture-word interference task: Effects of congruency and sequence manipulations"

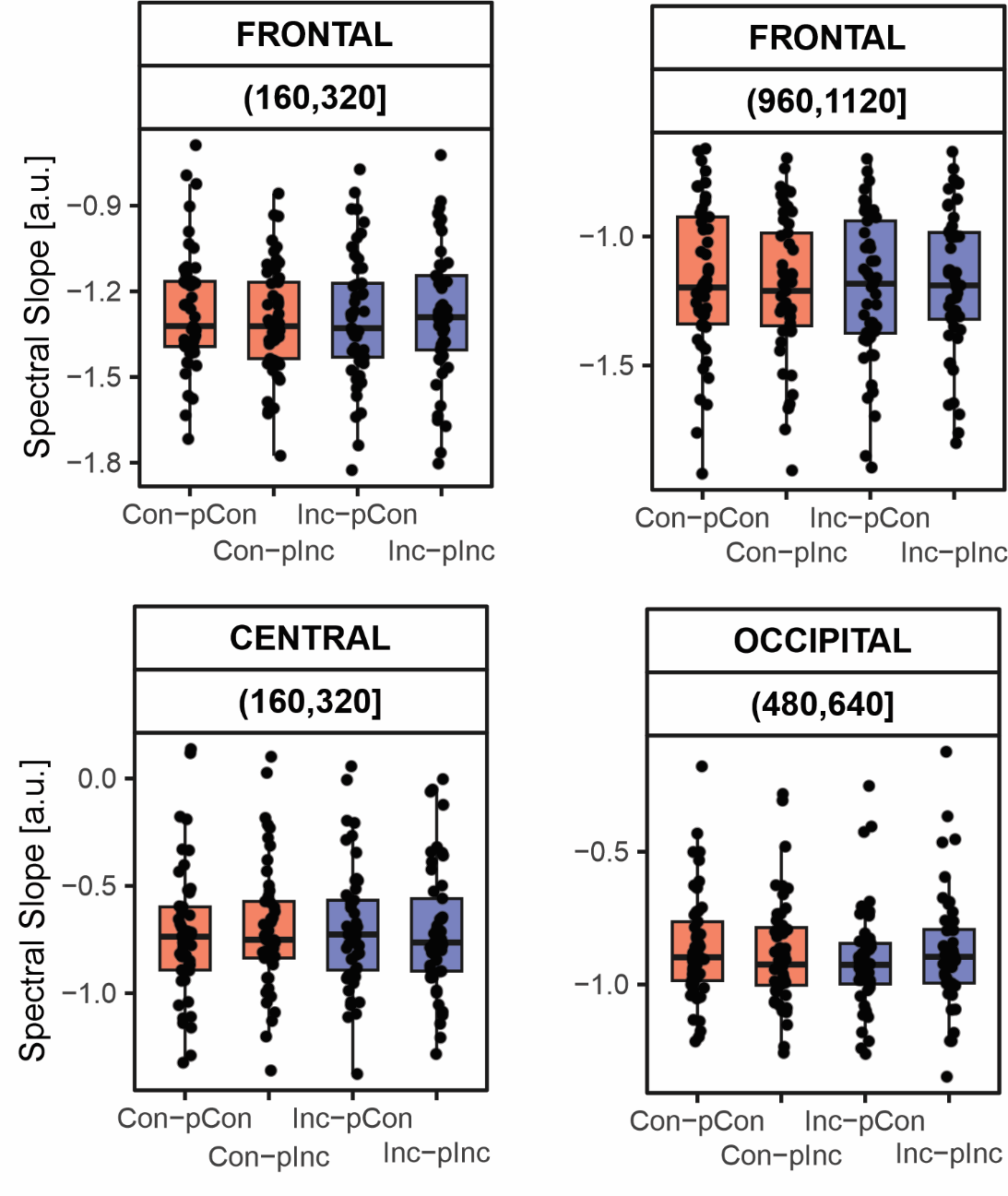

***Figure S1****.* **Current Congruency as a Function of Previous Congruency of Spectral Slope**. Boxplots of participant-level mean values illustrating current congruency as a function of previous congruency in the time windows and spatial components showing significant interactions in the repeated-measures, window-based exploratory ANOVA: 160–320 ms and 960–1120 ms in the frontal component (top left and top right), 160–320 ms in the central component (bottom left), and 480–640 ms in the occipital component (bottom right). Dots represent individual participant means, while boxplots summarize the distribution across participants. Condition labels are defined as follows: the first element (before the dash) indicates congruency in the current trial (Con = congruent, Inc = incongruent), and the second element (after the dash) indicates congruency in the preceding trial (pCon = previous congruent, pInc = previous incongruent).

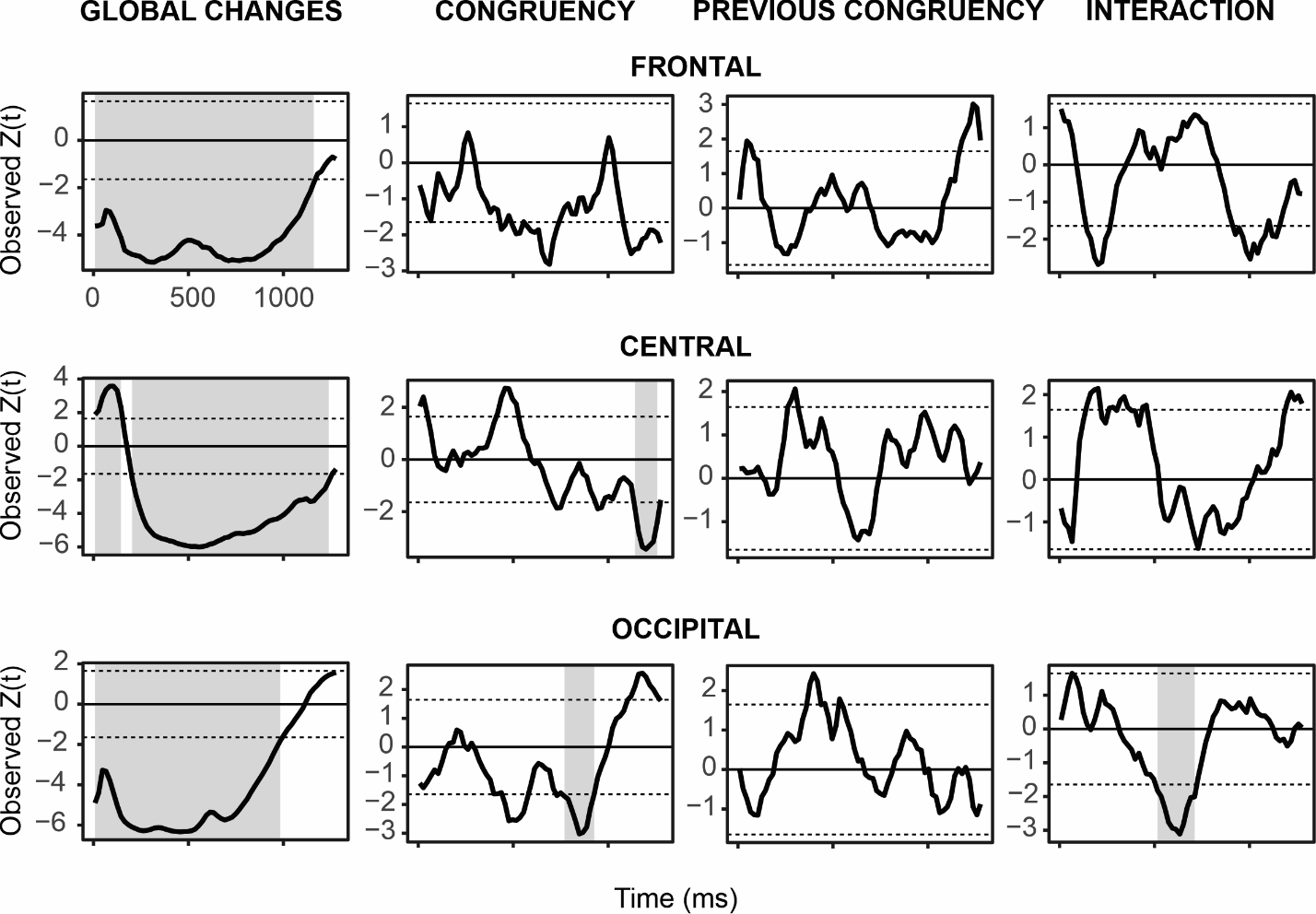

***Figure S2.* Time-Resolved Cluster-Based Permutation Results for Spectral Slope.** Panels display the *Z*-statistic time course (*Z*(*t*)) obtained from a sign-flip permutation procedure, with cluster formation thresholded at |*Z*| > 1.6449. Shaded regions indicate clusters significant at the family-wise error–corrected level (cluster-level *p_FWER_* < .05; max-intensity statistic, defined as the maximum |*Z*| within each cluster). From top to bottom: frontal, central, and occipital spatial components (PCA-derived scores). From left to right: global stimulus-induced deviations from the pre-stimulus baseline (baseline defined as the mean slope for *t* < 0 ms), followed by effects from the 2 × 2 design shown as within-subject contrasts: Congruency (Incongruent − Congruent), Previous Congruency (previous Incongruent − previous congruent), and their interaction, defined as the difference in the congruency effect as a function of Previous Congruency.

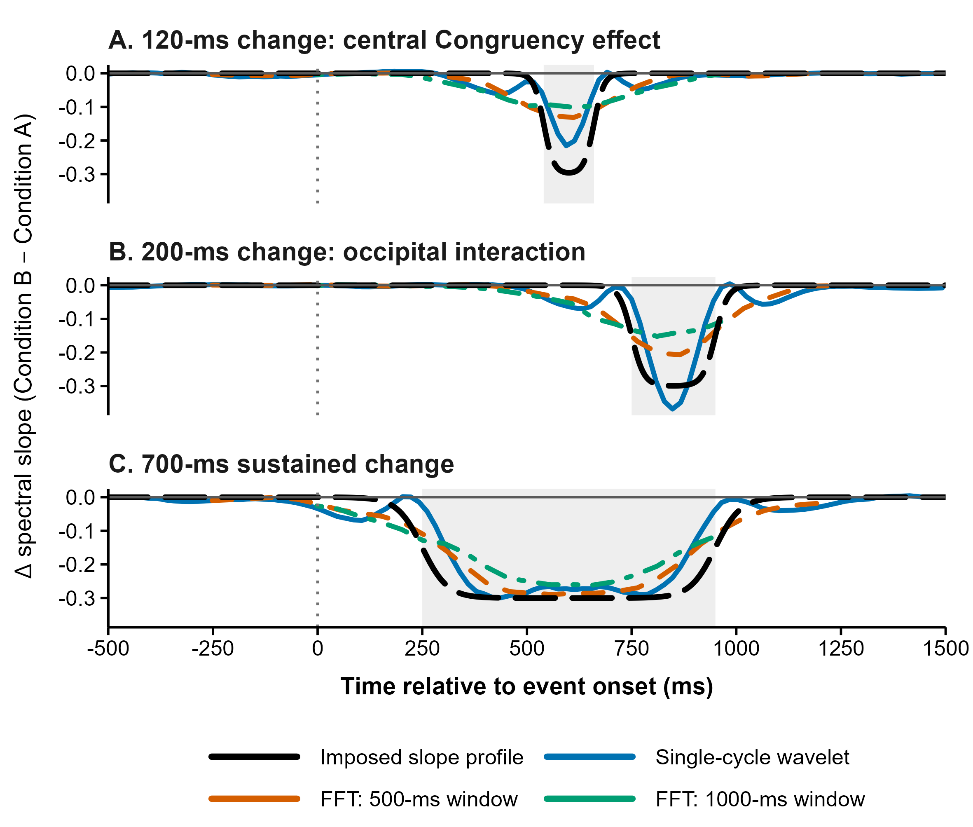

***Figure S3.* Sensitivity Analysis of Temporal Recovery. The figure compares the recovery of simulated aperiodic-slope changes using the single-cycle wavelet approach applied in the main analysis and conventional sliding FFT estimates based on fixed 500-ms and 1000-ms windows. The black dashed line shows the imposed slope profile, the blue line shows the single-cycle wavelet estimate, and the orange and green lines show the 500-ms and 1000-ms FFT estimates, respectively. Grey shading marks the interval during which the slope change was imposed. Paired synthetic EEG-like signals were generated with a baseline aperiodic slope of −1.20 in Condition A and an imposed slope change of −0.30 in Condition B. The two conditions shared Fourier phases and additive noise, allowing the comparison to isolate the imposed change in spectral slope. Three temporal profiles were simulated to approximate the durations of the key FWER-corrected effects reported in the manuscript: a 120-ms change corresponding to the late central Congruency effect observed from 1141 to 1258 ms, a 200-ms change corresponding to the occipital Congruency × Previous Congruency interaction observed from 516 to 711 ms, and a sustained 700-ms change representing the longer post-stimulus deviations from baseline. Each condition comprised 500 simulated trials. The simulated data were analysed using the same single-cycle wavelet procedure as in the main analysis and were compared with sliding FFT estimates based on fixed 500-ms and 1000-ms windows. For the wavelet analysis, power was estimated at 2.5, 5, 7.5, 15, and 25 Hz using one-cycle sine–cosine wavelet pairs, and spectral slopes were calculated from the log–log power–frequency relationship at successive time points. The FFT analyses used the same frequencies and estimated slopes within fixed windows advanced in 50-ms steps. Average event-related activity was removed separately within each condition before spectral estimation, matching the empirical analysis pipeline. All methods recovered the sustained 700-ms change. For the 120-ms and 200-ms changes, the fixed-window estimates were increasingly attenuated and temporally broadened, whereas the single-cycle estimator more closely followed the imposed temporal profile. The simulation evaluates recovery of the timing, direction, and approximate magnitude of an imposed broadband slope change. It is not intended to demonstrate that any estimator provides an unbiased or truly instantaneous estimate of the aperiodic slope.**
